# Plasma membrane and cytoplasmic compartmentalization: a dynamic structural framework required for pollen tube tip growth

**DOI:** 10.1101/2024.04.08.588543

**Authors:** Carolin Fritz, Theresa Maria Reimann, Jeremy Adler, Johanna Knab, Sylwia Schulmeister, Choy Kriechbaum, Sabine Müller, Ingela Parmryd, Benedikt Kost

## Abstract

Rapid, unidirectional pollen tube tip growth is essential for fertilization and is widely employed as a model of polar cell expansion, a process crucial for plant morphogenesis. Different proteins and lipids with key functions in the control of polar cell expansion are associated with distinct domains of the plasma membrane (PM) at the pollen tube tip. These domains need to be dynamically maintained during tip growth, which depends on massive secretory and endocytic membrane traffic. Very little is currently known about the regulatory and cellular mechanisms responsible for the compartmentalization of the pollen tube PM. To provide a reliable structural framework for the further characterization of these mechanisms, an integrated quantitative map was compiled of the relative positions in normally growing tobacco pollen tubes of PM domains 1) enriched in key signaling proteins or lipids, 2) displaying high membrane order, or 3) in contact with cytoplasmic structures playing important roles in apical membrane traffic. Previously identified secretory and endocytic PM domains were also included into this map. Internalization of regulatory proteins or lipids associated with PM regions overlapping with the endocytic domain was assessed based on brefeldin A (BFA) treatment. These analyses revealed remarkable aspects of the structural organization of tobacco pollen tube tips, which enhance our understanding of tip growth by providing important insights into 1) RAC/ROP signaling, 2) phosphatidylinositol 4,5-bisphosphate (PI4,5P_2_) metabolism and functions, 3) trafficking of signaling lipids, 4) functions of domains displaying high membrane order, and 5) Ca^2+^ regulation of secretion.

**Summary:** Quantitative mapping of plasma membrane and cytoplasmic domains at the tip of elongating tobacco pollen provides important insights into regulatory and cellular mechanisms essential for tip growth.

## Introduction

Pollen tubes, the male gametophytes of seed plants, deliver sperm cells enclosed in their vegetative cytoplasm to egg cells to enable fertilization (Bedinger, 1992; Dresselhaus et al. 2016). To exert this function, they elongate rapidly and strictly unidirectionally by tip growth, an extreme form of polarized cell expansion (Steer and Steer, 1989), which is also displayed e.g. by root hairs (Carol and Dolan, 2002) and apical cells of moss protonemata (Menand et al. 2007). In addition to mediating single cell morphogenesis, tip growth serves as an important model of directional plant cell expansion required for the development of multicellular organs (Kost et al. 2002; Orr et al. 2020). Pollen tube tip growth depends on massive secretion of cell wall material, which needs to be compensated by endocytic recycling of excess material delivered to the plasma membrane (PM) as a consequence of this process (Picton and Steer 1983; Derksen et al. 1995; Ketelaar et al. 2008). Conflicting models of the structural organization of apical membrane traffic underlying pollen tube tip growth have been proposed (Bove et al. 2008; Zonia and Munnik, 2008; Grebnev et al. 2017; Zhao et al. 2020). According to the well supported classical model, bulk secretion occurs apically and is balanced by massive lateral endocytic recycling of PM material (Picton and Steer, 1983; Derksen et al. 1995; Kost, 2008; Grebnev et al. 2020).

A cone-shaped cytoplasmic vesicle accumulation region (VAR) at the pollen tube apex is densely packed with secretory vesicles, which appear to be generated by a large subapical Trans-Golgi-Network (TGN) compartment. This TGN compartment encloses the VAR in the form of a torus and was proposed to integrate apical membrane traffic by also serving as recyling endosome (Stephan et al. 2014). Maintaining TGN positioning in a cytoplasmic region displaying rapid streaming depends on the myosin receptor NtRISAP and on the subapical cortical F-actin fringe (Stephan et al. 2014), a structure typically observed in pollen tubes (Kost et al. 1998; Lovy-Wheeler et al. 2005) with essential functions in tip growth (Vidali and Hepler, 2001; Hussey et al. 2006; Cai et al. 2015; Stephan, 2017). A steep tip-focused Ca^2+^ gradient generally detectable in the cytoplasm of tip-growing plant cells (Reiss and Herth, 1979; Tian et al. 2020) is thought to promote apical secretion and controls the direction of pollen tube growth (Malhó et al. 1994; Malhó and Trewavas, 1996). Although loss of this Ca^2+^ gradient has been shown to immediately arrest tip growth (Holdaway-Clarke and Hepler, 2003; Hepler et al. 2011; Michard et al. 2017), Ca^2+^ functions in this process are not well understood at the molecular level. Interestingly, coordinated oscillation of cell expansion, apical secretion and cytoplasmic Ca^2+^ concentration are frequently observed at the tip of elongating pollen tubes and root hairs (Holdaway-Clarke et al. 1999; Rounds et al. 2011).

RAC/ROP (RHO OF PLANT) GTPases typically associate with the PM due to posttranslational prenylation, specifically accumulate at the apical PM of cells undergoing tip growth, and function as key regulators of this process (Lin et al. 1996; Kost, 2008; Yalovsky et al. 2008; Guan et al. 2013; Scheible and McCubbin, 2019). These GTPases trigger downstream signaling when bound to GTP and are inactive in the GDP-bound conformation. RAC/ROP dependent signaling is controlled by different classes of upstream regulators. GAPs (GTPase ACTIVATING PROTEINs) inactivate RAC/ROP GTPases by promoting GTP hydrolysis (Klahre and Kost, 2006; Kost, 2010), whereas GEFs (GUANINE NUCLEOTIDE EXCHANGE FACTORs) do the opposite by stimulating GDP for GTP exchange (Berken et al. 2005; Gu et al. 2006). Both GAPs (Klahre and Kost 2006; Lauster et al. 2022; Ruan et al. 2023) and GEFs (Denninger et al. 2019; Le Bail et al. 2019) have been shown to associate with specific PM domains, although apart from REN subfamily GAPs they do not contain canonical membrane interaction domains or motives. GDIs (GUANINE NUCLEOTIDE DISSOCIATION INHIBITORs) act in concert with GDFs (GDI DISPLACEMENT FACTORs) to mediate RAC/ROP relocation between different PM domains. GDIs preferentially extract inactive RAC/ROP GTPases from the PM and form cytoplasmic heterodimers with them, which are destabilized by GDFs to promote RAC/ROP reassociation with the PM (Klahre et al. 2006; Ischebeck et al. 2011).

Membrane lipids also play important roles in the regulation of tip growth. The phosphoinositide phosphatidylinositol 4,5-bisphosphate (PI4,5P_2_) specifically accumulates in the PM at the apex of pollen tubes and root hairs (Kost et al. 1999; Kusano et al. 2008), where it was proposed to stimulate apical secretion and, at the same time, to act as GDF promoting RAC/ROP activation (Klahre et al. 2006; Kost, 2008). PM domains displaying high membrane order and the membrane lipid phosphatidylserine (PS) were reported to support nanoclustering specifically of active RAC/ROP GTPases at the PM of expanding root and leaf cells (Sorek et al. 2007, 2010; Platre et al. 2019; Pan et al. 2020), and may have the same function also in pollen tubes (Liu et al. 2009). Furthermore, pollen tube tip growth depends on specific accumulation of the signaling lipid phosphatidic acid (PA) within a lateral PM domain (Potocký et al. 2014; Pejchar et al. 2020; Scholz et al. 2022), although the molecular functions of this lipid in the control of tip growth remain to be clarified.

Remarkably, different regulatory proteins and lipids with important functions in the control of polar cell expansion maintain their association with distinct PM domains during pollen tube tip growth (Helling et al. 2006; Klahre and Kost, 2006; Sun et al. 2015; Stenzel et al. 2020; Scholz et al. 2022). Very little is currently known about the establishment and maintenance of these domains within the highly dynamic pollen tube PM, which is constantly remodeled by massive secretion and endocytic recycling. Evidently, the targeting of regulatory proteins and lipids to distinct domains of the pollen tube PM strongly depends on apical membrane traffic and vice versa. The further characterization of each of these two processes, as well as of their interplay, clearly is essential for a comprehensive understanding of tip growth. To provide a reliable structural framework for this challenging task, standardized procedures were employed to quantitatively determine the exact positions in normally growing tobacco pollen tubes of PM domains 1) enriched in key signaling proteins or lipids, 2) displaying high membrane order, or 3) in direct contact with cytoplasmic structures or regions playing important roles in apical membrane traffic. A quantitative map of the PM at the tip of tobacco pollen tubes was compiled, which integrates functional PM regions characterized here with previously identified apical secretory and lateral endocytic domains (Grebnev et al. 2020). In addition, brefeldin A (BFA) treatment was employed to assess possible endocytic recycling of regulatory proteins or lipids associated with PM regions that are overlapping with the lateral endocytic PM domain.

## Results and discussion

### Quantitative structural analysis of growing tobacco pollen tube tips

Our study builds on and integrates a previously reported quantitative model of membrane traffic and PM compartmentalization at the tip of tobacco pollen tubes developed by Grebnev et al. (2020). These authors have investigated steady-state distributions as well as dynamics of FM4-64-labeled lipids and of YFP-tagged transmembrane (TM) proteins in normally growing or BFA-treated tobacco pollen tubes to quantitatively characterize the spatial organization of bulk membrane traffic underlying tip growth. Based on these analyses, bulk secretion was demonstrated to deliver lipids and transmembrane proteins to an apical PM domain extending from 0 - 3.5 µm meridional distance (MD, measured along the curved PM) from the apex (Fig. 1; secretory domain [green]). Apical secretion was revealed to be compensated by massive endocytic recycling of membrane lipids within a lateral PM domain extending from 5.9 - 14.8 µm MD from the apex (Fig. 1; endocytic domain [green]), which was associated with AtAP180, a marker for clathrin-mediated endocytosis (Barth and Holstein, 2004; Kaneda et al. 2019). Importantly, bulk fusion of secretory vesicles with the apical secretory domain together with massive membrane internalization within the lateral endocytic domain were proposed to result in constant retrograde drift of membrane components between these two PM regions (Fig. 1; black arrowheads). Furthermore, Grebnev et al. (2020) found that the previously discovered cortical F-actin fringe (Kost et al. 1998; Lovy-Wheeler et al. 2005) is in contact with the PM exactly between the apical secretory and the lateral endocytic domains (Fig. 1; fringe [green]), where it partially overlaps with a subapical TGN compartment required for membrane recycling (Fig. 1; TGN [green]). Consistent with this observation, subapical TGN positioning depends on the F-actin fringe (Stephan et al. 2014).

**Figure 1:**
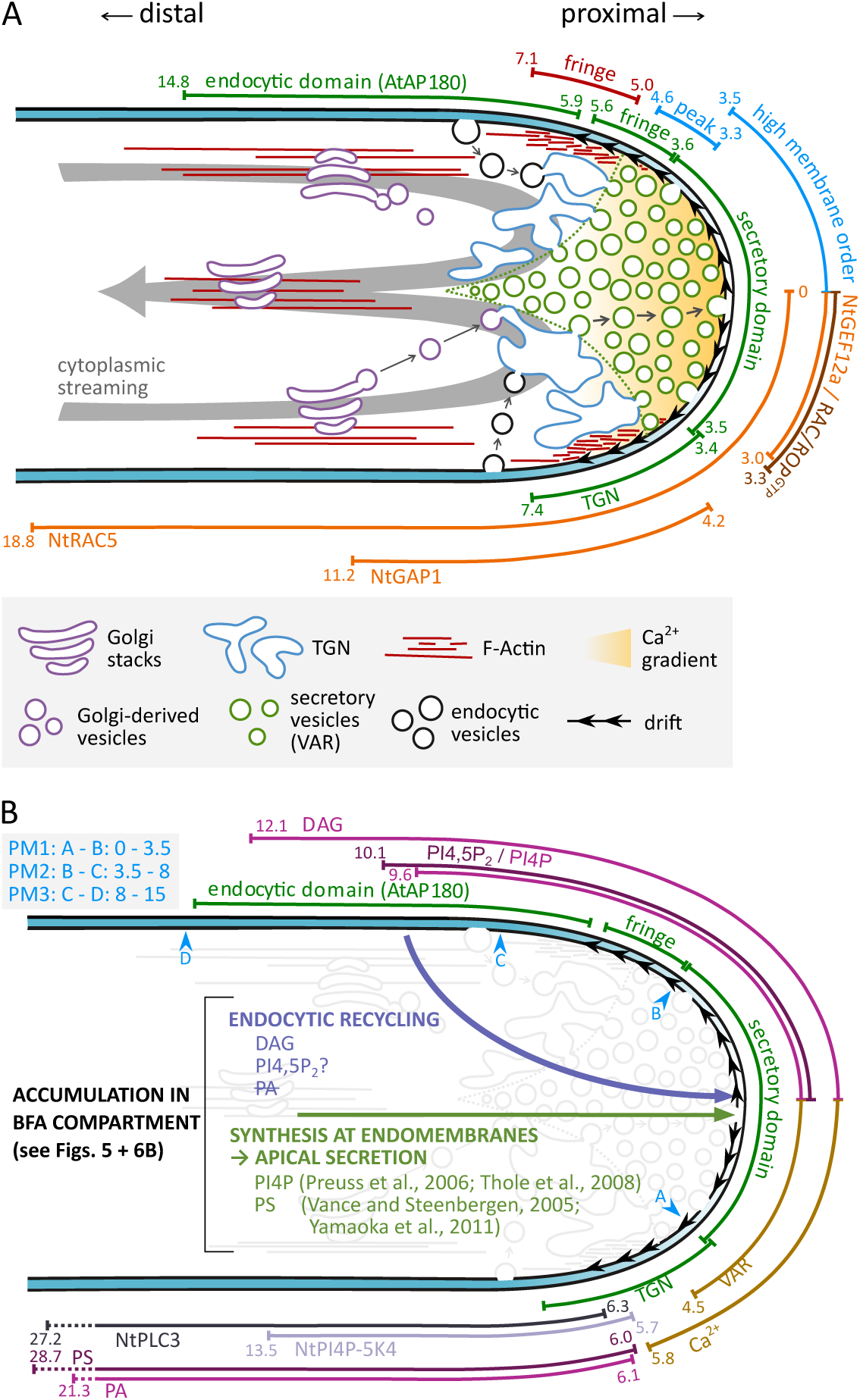
Quantitative structural organization of growing tobacco pollen tube tips. Integrated model of cytoplasmic organization, apical membrane traffic and PM compartmentalization based on data reported by Grebnev et al. (2020) and resulting from quantitative analyses presented here. PM domains identified by Grebnev et al. (2020) are indicated in green. Fringe and TGN domains: PM regions in contact with the cortical F-actin fringe and the subapical Trans-Golgi-Network compartment, respectively. Peak: membrane order peak. PM1 - 3: PM regions selected for the determination of average membrane order. Numbers represent meridional distances (MDs) from the apex (measured along the curved PM) reported by Grebnev et al. (2020) or determined based on quantitative analyses described here (Figs. 3 and 7A). PM domains are drawn to scale. Proximal to distal: increasing distance from the apex. **A)** Model of apical membrane traffic proposed by Grebnev et al. (2020): a) proteins and lipids delivered to the secretory PM domain by apical exocytosis are recycled within the lateral endocytic domain, resulting in retrograde drift within the apical PM (black arrowheads), b) apical membrane traffic is integrated by the subapical TGN compartment, which repacks proteins and lipids delivered to its distal surface by Golgi-derived or endocytic vesicles into secretory vesicles that are released from its proximal surface, and c) stable positioning of the subapical TGN compartment within a rapidly streaming cytoplasmic region depends on the cortical F-actin fringe. Quantitative mapping described here enabled incorporation into this model of PM domains associated with RAC/ROP signaling proteins (orange/brown) or displaying high membrane order (blue), which appear to have important, possibly interdependent functions in the regulation of apical membrane traffic. Furthermore, data presented here establish striking overlap between apical cytoplasmic regions densely packed with secretory vesicles (vesicle accumulation region: VAR) or displaying elevated cytoplasmic Ca^2+^ concentrations (orange shading). These data also place the PM region in contact with the cortical F-actin fringe (red) 1.4 - 1.5 µm further away from the apex than previously observed (green; Grebnev et al. 2020). **B)** Complementation of the model presented in A) with PM domains quantitatively mapped here, which a) are associated with specific lipids (pink/purple) or lipid-modifying enzymes (grey) with important functions in the regulation of tip growth, or b) are in direct contact with the VAR or with a cytoplasmic region displaying elevated cytoplasmic Ca^2+^ concentrations (brown). Text superimposed onto the pollen tube cytoplasm provides information concerning the biosynthesis and trafficking of analyzed lipids, which is supported by the indicated references as well as by brefeldin A (BFA) effects on the intracellular distribution of these lipids characterized here (Figs. 5 and 6B).

To extend the model described in the previous paragraph, additional PM and cytoplasmic domains, which are known or believed to have important regulatory or structural functions during tip growth, were quantitatively mapped (Fig. 1). To this end, the intracellular distributions of eYFP-tagged tobacco pollen tube RAC/ROP signaling proteins, and of fluorescent fusion proteins serving as makers for lipids, Ca^2+^, organelles or F-actin, were quantitatively analyzed in stably transformed (with one exception as indicated below) tobacco pollen tubes, which elongated at normal rates (≥ 3.0 µm/min, Supplemental Fig. S1; Grebnev et al. 2020) and displayed no morphological defects. Furthermore, normally growing tobacco pollen tubes (Supplemental Fig. S1) labeled with Di-4-ANEPPDHQ were employed to map variations in membrane order, which determine physical membrane characteristics and regulate membrane-associated signaling (Ashrafzadeh and Parmryd 2015; Sezgin et al. 2017). To enable comparative analyses, standardized procedures were employed a) to quantify marker protein fluorescence at or directly underneath the PM (PM-associated relative fluorescence intensity [RFI]) at different MDs from the apex, and b) to determine the MD from the apex of the proximal (closer to the apex) and distal (further away from the apex) ends of all investigated PM domains.

### PM association patterns of RAC/ROP signaling proteins shed light on RAC/ROP^GTP^ polarization and functions

NtRAC5 (Kieffer et al. 2000) is a pollen tube-specific tobacco RAC/ROP GTPase, which plays a key role in the control of tip growth (Klahre et al. 2006; Kost, 2008). Moderate overexpression of this protein effectively depolarizes tobacco pollen tube tip growth and results in massive apical ballooning (Klahre et al. 2006). Imaging the intracellular distribution of fluorescent NtRAC5 fusion proteins in normally elongating tobacco pollen tubes represents a substantial challenge, as expressing these fusion proteins even at relatively low levels strongly affects pollen tube growth and morphology (Sun et al. 2015). However, quantitative analysis of transiently transformed tobacco pollen tubes expressing an eYFP-NtRAC5 fusion protein at the lowest level detectable by confocal microscopy established that during normal tip growth this fusion protein specifically associates with the PM within an apical domain extending from 0 - ca. 20 µm MD from the apex (Sun et al. 2015). To re-evaluate these data, transgenic tobacco lines were generated, which expressed the same eYFP-NtRAC5 fusion protein under the control of the pollen-specific *Lat52* promoter (Twell et al. 1990). Many of these lines displayed severely disrupted pollen tube growth resulting from eYFP-NtRAC5 overexpression, along with strong unspecific association of this fusion protein with the entire pollen tube PM. However, several transgenic lines expressing eYFP-NtRAC5 at the lowest detectable level were also identified. Pollen tubes of these lines consistently displayed normal growth (Supplemental Fig. S1) and weak eYFP-NtRAC5 association specifically with an apical PM domain (Fig. 2A), although the extension of this domain varied considerably between individual pollen tubes with identical genetic background (Supplemental Fig. S2). Quantitative analysis of such pollen tubes showed that starting at a MD of about 20 µm from the apex the RFI representing eYFP-NtRAC5 association with the PM gradually increased towards the extreme tip (Fig. 2A), which is consistent with the previously reported data (Sun et al. 2015). In control experiments, accumulation of free eYFP at the PM was not detected (Fig. 2A).

**Figure 2:**
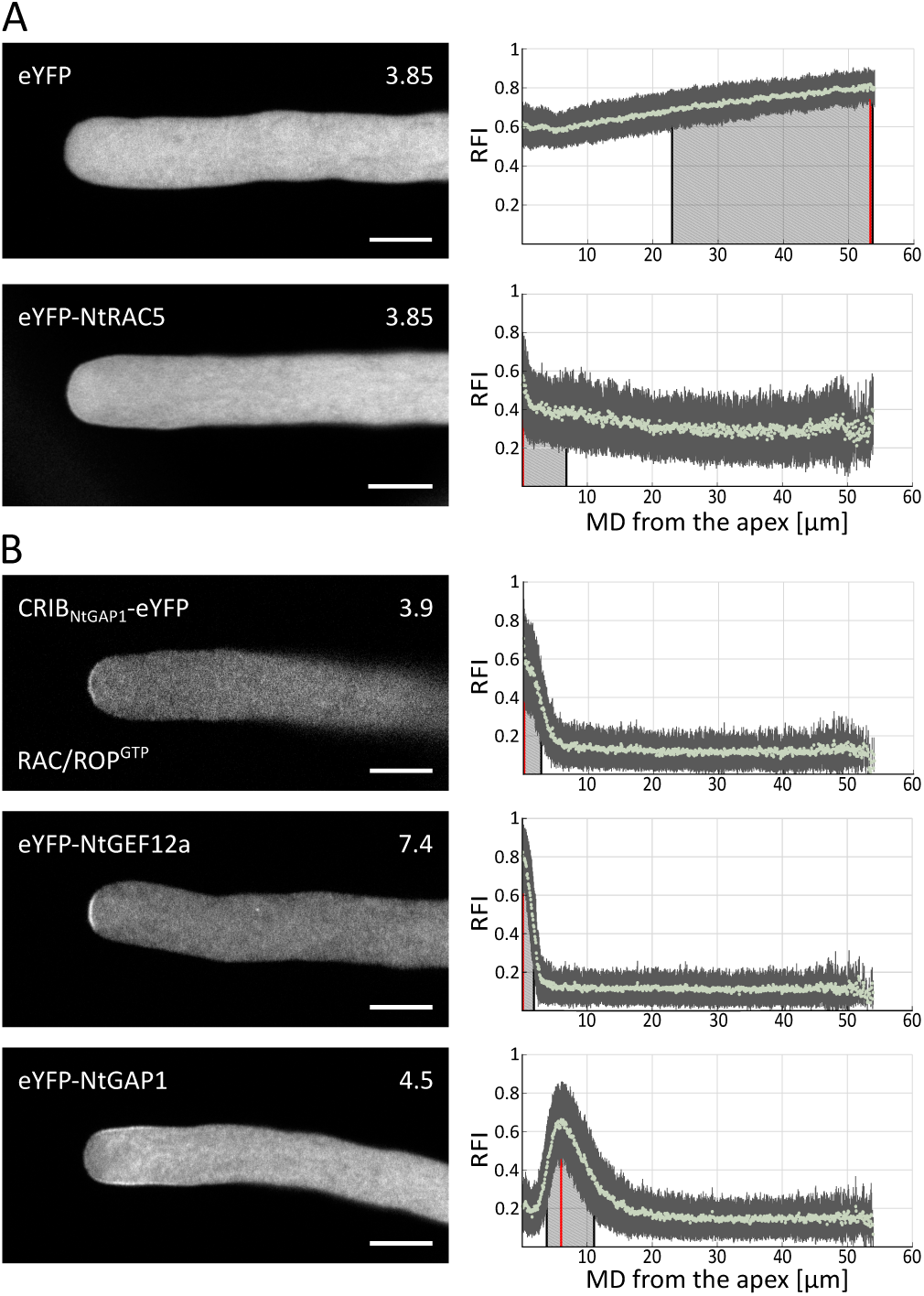
PM association of RAC/ROP signaling proteins. **Left column:** medial confocal optical sections through representative pollen tubes expressing the indicated eYFP fusion proteins, or free eYFP used as control, either transiently (CRIB_NtGAP1_-eYFP) or stably (all other fluorescent proteins). Numbers (top right) indicate the growth rate (µm/min) of the individual pollen tubes shown, which was determined after image acquisition (average growth rate of all analyzed pollen tubes: Supplemental Fig. S1). Scale bars: 8 µm. **Right column:** quantitative analysis of PM-associated fluorescence (directly underneath [eYFP] or at the PM) in all analyzed pollen tubes expressing each indicated protein. Light green dots: mean relative fluorescence intensity (RFI) associated with the PM at different meridional distances (MDs) from the apex (MD = 0 µm). Both sides of imaged pollen tubes were analyzed, if possible. Dark grey vertical lines: standard deviation. Red vertical lines: maximal RFI. Black vertical lines delimiting light grey shading: proximal and distal half-maximal RFI (if applicable). **A)** Intracellular distribution and PM association of eYFP-NtRAC5 (n = 49 pollen tubes, 95 RFI distribution patterns), along with the intracellular distribution of free eYFP (n = 55 pollen tubes, 110 RFI distribution patterns). **B)** Intracellular distribution and PM association of CRIB_NtGAP1_-eYFP serving as a marker for GTP-bound active RAC/ROP^GTP^ (n = 61 pollen tubes, 118 RFI distribution patterns), as well as of eYFP fused to NtGEF12a (n = 55 pollen tubes, 110 RFI distribution patterns) or to NtGAP1 (n = 65 pollen tubes, 128 RFI distribution patterns), which activate or inactivate RAC/ROP signaling functions, respectively.

The apical NtRAC5 domain shown in figure 2A is expected to contain active NtRAC5^GTP^ along with inactive NtRAC5^GDP^. As previously reported (Klahre and Kost, 2006), eYFP fused to the isolated CRIB (CDC42/RAC INTERACTIVE BINDING) domain of NtRhoGAP1 (CRIB_NtGAP1_-eYFP), which serves as a specific maker for active RAC/ROP^GTP^ (Wu et al. 2000, 2001), strongly and specifically associates with a small PM domain at the pollen tube apex (Fig. 2B). Because transgenic CRIB_NtGAP1_-eYFP expressing lines could not be established, transient expression was employed to further characterize the intracellular distribution of this fusion protein. Quantitative analysis (Fig. 2B) confirmed that the RAC/ROP^GTP^ domain detected by CRIB_NtGAP1_-eYFP only overlapped with a small apical region of the much larger NtRAC5 domain (Fig. 2A), suggesting that the remainder of the NtRAC5 domain contains NtRAC5^GDP^. Importantly, the RAC/ROP^GTP^ domain was nearly identical to the apical secretory domain identified by Grebnev et al. (2020; Fig. 1A), supporting a key role of active NtRAC5^GTP^, and possibly other RAC/ROP^GTP^ proteins, in promoting the fusion of secretory vesicles with the PM.

AtROPGEF12 was reported to play a crucial role in RAC/ROP activation at the tip of Arabidopsis pollen tubes downstream of TM receptors of the RLK (RECEPTOR-LIKE KINASE) family (Kaothien et al. 2005; Zhang and McCormick, 2007). Screening available tobacco genome databases resulted in the identification of three full-length genes encoding close AtROPGEF12 homologs, which were designated NtGEF12a, b and c (Supplemental Fig. S3A - C). Like their Arabidopsis homolog, all three NtGEF12 isoforms a) are specifically expressed in pollen tubes (Supplemental Fig. S3D and E), b) comprise a conserved central PRONE (PLANT SPECIFIC ROP NUCLEOTIDE EXCHANGER) domain (Supplemental Fig. S3A) that can mediate RAC/ROP activation by promoting GDP for GTP exchange (Berken et al. 2005), and c) contain three conserved S/T residues near the C-terminus (Supplemental Fig. S3A), whose RLK-mediated phosphorylation was proposed to relieve autoinhibition of nucleotide exchange activity (Zhang and McCormick, 2007).

Interestingly, eYFP fused to NtGEF12a, which shares the highest amino acid identity with AtROPGEF12 (Supplemental Fig. S3B and C) and interacts with NtRAC5 in split-ubiquitin yeast two hybrid assays (Supplemental Fig. S4), strongly associated with the pollen tube PM specifically within an apical domain that almost perfectly overlapped with the RAC/ROP^GTP^ domain (Fig. 2B). This observation supports a key function of NtGEF12a in RAC/ROP activation at the pollen tube apex, which appears to promote apical secretion as discussed above. In apical protonemal cells of the moss *Physcomitrium patens*, which also expand by tip growth, PpROP1 and PpROPGEF4 expressed at endogenous levels are associated with very similar apical PM domains as NtRAC5 and NtGEF12a, respectively, in tobacco pollen tubes (Le Bail et al. 2019; Cheng et al. 2020). This indicates that molecular mechanisms playing essential roles in the control of apical RAC/ROP activity and secretion during tip growth may be conserved among land plants.

As previously reported (Klahre and Kost, 2006), eYFP-tagged NtRhoGAP1 (henceforth abbreviated NtGAP1), which inactivates RAC/ROPs by promoting GTP hydrolysis, specifically associates with a lateral domain of the pollen tube PM (Fig. 2B). Quantitative analysis (Fig. 2B) established that PM labeling by this fusion protein bordered on the apical RAC/ROP^GTP^ domain and substantially overlapped with the presumably NtRAC5^GDP^-associated lateral part of the NtRAC5 domain (Fig. 1A). These observations strongly support a previously proposed model (Klahre and Kost, 2006; Kost, 2008) suggesting that NtGAP1 contributes to polarity maintenance by laterally promoting RAC/ROP GTPase activity and by thereby restricting RAC/ROP^GTP^ spreading, which is driven by retrograde drift within the apical PM. This model also proposed that maintenance of apical RAC/ROP^GTP^ polarization during pollen tube tip growth further requires constant GDI-mediated recycling of inactive RAC/ROP^GDP^ from the lateral to the apical PM, which is followed by GEF-mediated RAC/ROP reactivation. This concept is perfectly consistent with the apical colocalization of the NtGEF12a and RAC/ROP^GTP^ domains shown here (Figs. 1A and 2B).

To further enhance our understanding of the control of pollen tube tip growth, it will be essential to characterize the molecular and cellular mechanism responsible for the correct targeting of NtGEF12a and NtGAP1 to their respective PM domains. Key aims of this research will be the identification a) of RLKs that activate NtGEF12a and possibly contribute to the recruitment of this protein to the apical PM, as well as b) of interaction partners binding to the N-terminal NtGAP1 domain, which was demonstrated to be responsible for the specific targeting of this protein to the lateral PM (Klahre and Kost, 2006).

### Membrane order is high at the apex and shows a peak at its flanks

Dynamic heterogeneity within the PM, which includes local variation in membrane order, modulates physical membrane characteristics and plays important roles in the regulation of PM-associated signaling. Embedded within regions exhibiting low membrane order (liquid-disordered regions), the PM contains transiently formed nanodomains displaying high membrane order (liquid-ordered regions), a liquid crystalline state characterized by regular arrangement and dense packing of hydrophobic tails within lipid bilayers. Nanodomains displaying high membrane order, often referred to as “lipid rafts”, are typically enriched in sterols, sphingolipids and saturated phospholipids, which are thought to interact with each other and with specific proteins to establish signaling platforms (Sezgin et al. 2017).

Membrane order and anionic lipids such as phosphatidylserine (PS) promote nanoclustering of animal RAS and RHO family small GTPases at the PM, which controls the signaling activity of these proteins (Sartorel et al. 2018; Van et al. 2021). Nanoclustering of plant RAC/ROP GTPases at the PM also appears to regulate signaling triggered by these proteins. Activation-dependent S-acylation of the Arabidopsis RAC/ROP GTPase AtROP6 causes this protein to accumulate in lipid raft-enriched “detergent resistant membrane (DRM)” fractions. Point mutations preventing S-acylation not only abolished DRM partitioning of AtROP6, but also disrupted the signaling activity of this protein (Sorek et al. 2007, 2010). Recently reported microscopic investigations showed that auxin, a plant hormon and growth factor, induces AtROP6 nanoclustering at the PM of root and leaf epidermal cells in a PS- or sterol-dependent manner, respectively (Platre et al. 2019; Pan et al. 2020). Auxin-induced sterol-dependent AtROP6 nanoclustering was proposed to play an important role in promoting lobe expansion during the development of interdigitating leaf epidermal pavement cells. Consistent with this hypothesis, high membrane order within the PM of these cells was specifically detected at the tip of growing lobes based on labeling with Di-4-ANEPPDHQ (Pan et al. 2020), a fluorescent dye whose emission spectrum is shifted towards shorter wave lengths in PM regions enriched in liquid-ordered domains, as compared to PM regions containing larger proportions of liquid-disorderd domains (Jin et al. 2006).

Interestingly, Di-4-ANEPPDHQ labeling of the PM also indicated high membrane order at the tip of growing gymnosperm pollen tubes (Liu et al. 2009), and large scale proteomics established sterol-dependent DRM partitioning of RAC/ROP GTPases and other RAC/ROP signaling proteins, which are expressed in rice pollen (Han et al. 2018). Furthermore, point mutations disrupting conserved S-acylation sites reduced the ability of NtRAC5 to depolarize tobacco pollen tube growth upon overexpression (Sun et al. 2015), suggesting that similar to AtROP6 this protein may require S-acylation-dependent lipid raft association to effectively stimulate downstream signaling. To further investigate this possibility, Di-4-ANEPPDHQ labeling was employed to identify and map regions of the tobacco pollen tube PM displaying high membrane order.

Based on widefield imaging of Di-4-ANEPPDHQ fluorescence, average generalized polarization (GP), an indicator of membrane order (Jin et al. 2006; Parasassi et al. 1990; Zhao et al. 2015), was determined in three adjacent PM regions at the tip of normally growing (Supplemental Fig. S1) pollen tubes: a) PM1 (0 - 3.5 µm MD from the apex) corresponding to the nearly identical apical NtGEF12a, RAC/ROP^GTP^ and secretory domains, b) PM 2 (3.5 - 8 µm MD from the apex) largely overlapping with the PM contact sites of the cortical F-actin fringe and the subapical TGN compartment, and c) PM3 (8 - 15 µm MD from the apex) covering most of the lateral endocytic domain (Figs. 3A and 1B). Interestingly, the average GP-value was substantially higher in PM1 than in PM2 and PM3 (Fig. 3B), indicating enrichment of PM1 in lipid rafts that evidently are not individually resolved by Di-4-ANEPPDHQ imaging. High membrane order specifically within the apical RAC/ROP^GTP^ and secretory domain suggests that lipid raft partitioning may promote the ability of active NtRAC5^GTP^ to stimulate secretion, as proposed in the previous paragraph.

**Figure 3:**
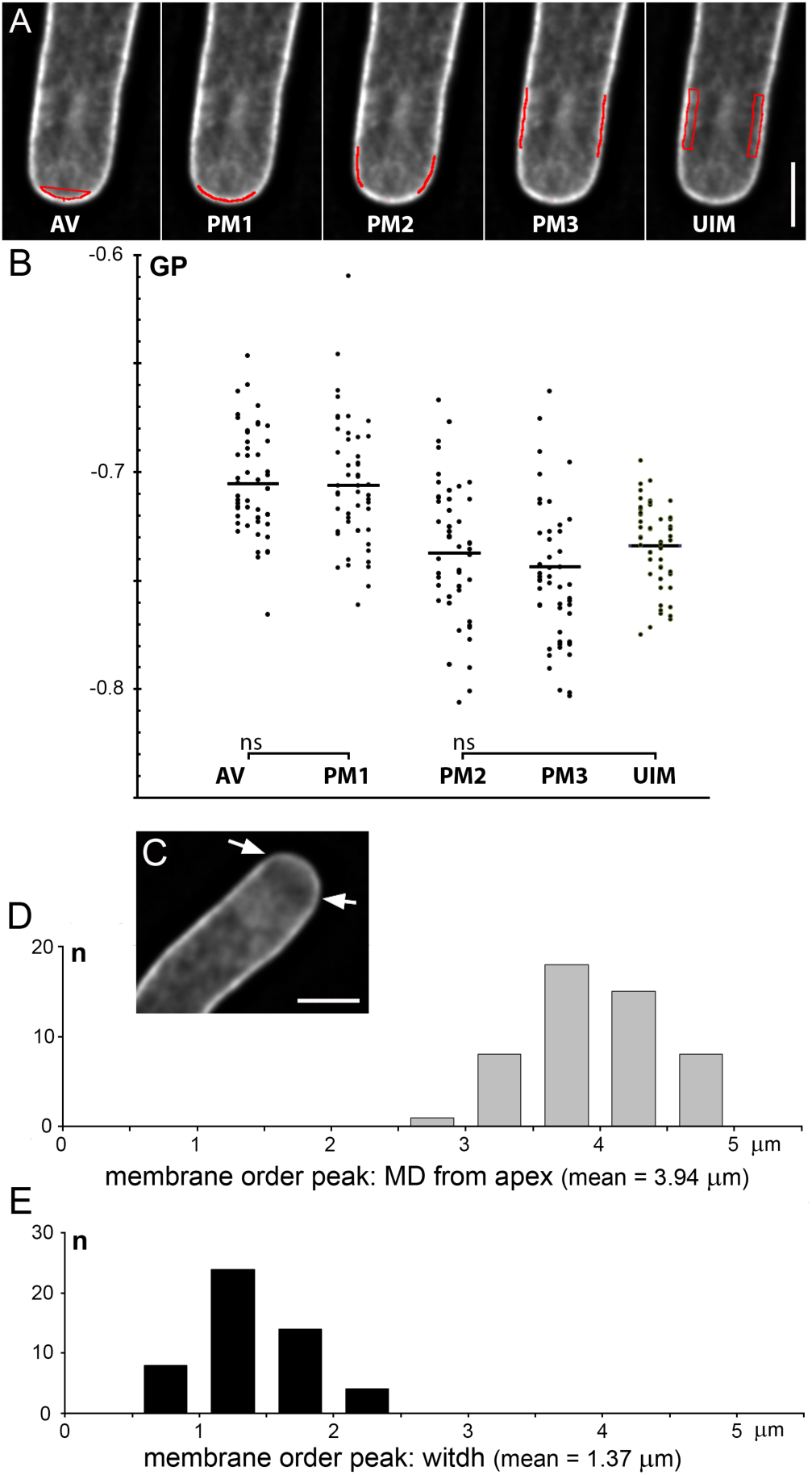
Membrane order within different PM domains and cytoplasmic regions containing intracellular membranes. Normally growing pollen tubes (Supplemental Fig. S1) stained with Di-4-ANEPPDHQ were imaged by widefield fluorescence microscopy. Z-stacks around the medial pollen tube plane were simultaneously acquired in two channels (510 - 550 nm, ≥ 645 nm) and deconvolved. Resulting image pairs were employed to determine generalized polarization (GP), which indicates membrane order (i.e. the relative proportion of liquid-ordered and liquid-disordered domains). **A)** Red lines or shapes overlaid onto an image of a Di-4-ANEPPDHQ-stained pollen tube indicate regions of interest (ROIs), within which average GP-values were determined. Analyzed ROIs covered three distinct PM domains (PM1 - 3), and two cytoplasmic regions containing either exclusively apical vesicles (AV, underneath PM1) or unspecified intracellular membranes enclosing different cytoplasmic organelles (UIM, underneath PM3). Meridional distances (MDs) from the apex spanned by the PM ROIs were 0 - 3.5 µm (PM1), 3.5 - 8 µm (PM2) or 8 - 15 µm (PM3). Scale bar: 8 µm. **B)** Average GP-values within the different ROIs defined in A) determined for individual pollen tubes (dots), along with the means of these values (horizontal lines). GP-values closer to 0 indicate higher membrane order (i.e. a larger proportion of liquid-ordered domains). n = 50. When image alignment allowed, the two ROIs on each side of analyzed pollen tubes were combined to determine individual average PM2, PM3 and UIM GP-values (36 of 50 pollen tubes analyzed). Data were statistically analyzed using pairwise two-tailed t-test. ns (not significantly different): p > 0.05. Pairwise analysis of all other data sets indicated significant differences: p ≤ 0.001. **C)** Representative deconvolved image of a Di-4-ANEPPDHQ-stained pollen tube acquired in the upper (≥ 645 nm) channel showing a dip in fluorescence intensity roughly at the border between PM1 and PM2 (arrows), which indicates a peak in membrane order at this location. Scale bar: 8 µm. **D)**, **E)** Investigation of GP-value distribution along the PM confirmed the presence of a peak in membrane order at the location indicated in C) in all pollen tubes analyzed. n = 50. Histograms show the number of pollen tubes (Y-axis) in which the membrane order peak displayed a MD from the apex (D), or a width (E), within the indicated range. When measurements were possible on both sides of analyzed pollen tubes, the means of the two obtained values were used as individual data points (36 of 50 pollen tubes analyzed). The membrane order peak displayed a mean MD from the apex of 3.94 µm [95 % confidence interval: ± 0.20] and a mean width of 1.37 µm [95 % confidence interval: ± 0.17].

GP-analysis of Di-4-ANEPPDHQ imaging data further demonstrated that PM1 and a cytoplasmic region enriched in apical vesicles (AV) located directly below this PM domain display equally high membrane order (Fig. 3A and B). Interestingly, the same significantly lower membrane order was detected in PM3 and in a cytoplasmic region directly underneath this PM domain (Fig. 3A and B), which contained unspecified intracellular membranes (UIM) enclosing different organelles such as ER, Golgi, mitochondria, plastids, peroxisomes, and endocytic vesicles (Cheung and Wu, 2007). These observations are consistent with the model of apical membrane traffic displayed in figure 1A, and indicate that a) high membrane order may be established in PM1 upon fusion of apical vesicles with this domain, and b) lower membrane order displayed by PM3 may be maintained in endocytic vesicles pinching off this PM region.

Interestingly, imaging Di-4-ANEPPDHQ emission at wave lengths longer than 645 nm, which is reduced in membrane regions displaying high membrane order (Jin et al. 2006), indicated the existence of a peak in membrane order roughly at the border between the apical PM1 and the lateral PM2 domains (Fig. 3C). Examination of GP-value distribution in this border region confirmed the consistent presence of a local peak in membrane order in the PM of all analyzed pollen tubes, which displayed an average MD from the apex of 3.9 µm (Fig. 3D) and a mean width of 1.4 µm (Fig. 3E). This peak overlaps with the distal end of the apical domain displaying high membrane order and with the proximal end of the cortical F-actin fringe (Fig. 1A). As discussed in more detail below, the observed lateral peak in membrane order may play a role in maintaining structural organization at the pollen tube tip, possibly depending on functional interactions with the F-actin fringe. Supporting this hypothesis, cortical actin filaments in T cells have been shown to induce the formation of highly ordered membrane domains upon association with the PM (Dinic et al. 2013).

To further investigate structural and regulatory functions of PM regions displaying high membrane order in tobacco pollen tubes, DRM-associated proteins purified from pollen tube extracts need to be identified and their intracellular distribution during normal tip growth must be determined using high-resolution fluorescence microscopy. It will be particularly interesting to investigate DRM partitioning of NtRAC5 as well as possible nanoclustering of this protein within the apical RAC/ROP^GTP^ domain. Exploring NtRAC5 nanoclustering along the curved surface of the apical dome of growing pollen tubes, of which high-resolution images perpendicular to the longitudinal axis are hard to obtain, will depend on technical innovation possibly including the adaptation of light sheet microscopy. TIRF-based imaging methods require membranes to be imaged in the immediate proximity of the cover slip and are therefore not readily applicable for this purpose, although they have been instrumental for analyses of AtROP6 nanoclustering reported in the literature (Platre et al. 2019; Pan et al. 2020).

### PM regions involved in phosphoinositide signaling provide insights into the formation and the functions of an apical PI4,5P_2_ domain

Phosphatidylinositol 4,5-bisphosphate (PI4,5P_2_) is an important signaling lipid with diverse functions in eukaryotic cells (Meijer and Munnik, 2003; Heilmann, 2016; Katan and Cockcroft, 2020; Li et al. 2020). This lipid accumulates in the PM specifically at the pollen tube tip (Kost et al. 1999; Ischebeck et al. 2008), where it was proposed to act both as a downstream effector and an upstream regulator of RAC/ROP activity (Kost, 2008). Pollen tube RAC/ROP GTPases physically interact with phosphatidylinositol 4-phosphate 5-kinases (PI4P-5Ks; Kost et al. 1999; Fratini et al. 2021), which phosphorylate phosphatidylinositol 4-phosphate (PI4P) to generate PI4,5P_2_ (Fig. 4A), supporting a role of PI4,5P_2_ in RAC/ROP downstream signaling. Acting as a RAC/ROP effector, PI4,5P_2_ may promote polarized secretion required for pollen tube tip growth, as it does during related cellular processes (Hay et al. 1995). Consistent with this hypothesis, enhanced PI4,5P_2_ production resulting from PI4P-5K overexpression results in massive deposition of cell wall material at the pollen tube tip, which is mediated by apical secretion (Ischebeck et al. 2008; Sousa et al. 2008; Fratini et al. 2021). Furthermore, both in mammalian cells (Fauré et al. 1999; DerMardirossian and Bokoch, 2005) and in tobacco pollen tubes (Ischebeck et al. 2011; Fratini et al. 2021) PI4,5P_2_ was reported to display GDF activity, which promotes PM association and subsequent GEF-mediated activation of RHO GTPases. Positive feedback between RAC/ROP-stimulated PI4,5P_2_ production and PI4,5P_2_-mediated RAC/ROP activation potentially promotes apical polarization of RAC/ROP signaling and of cell expansion at the pollen tube tip (Kost et al. 1999; Kost, 2008; Ischebeck et al. 2011).

**Figure 4:**
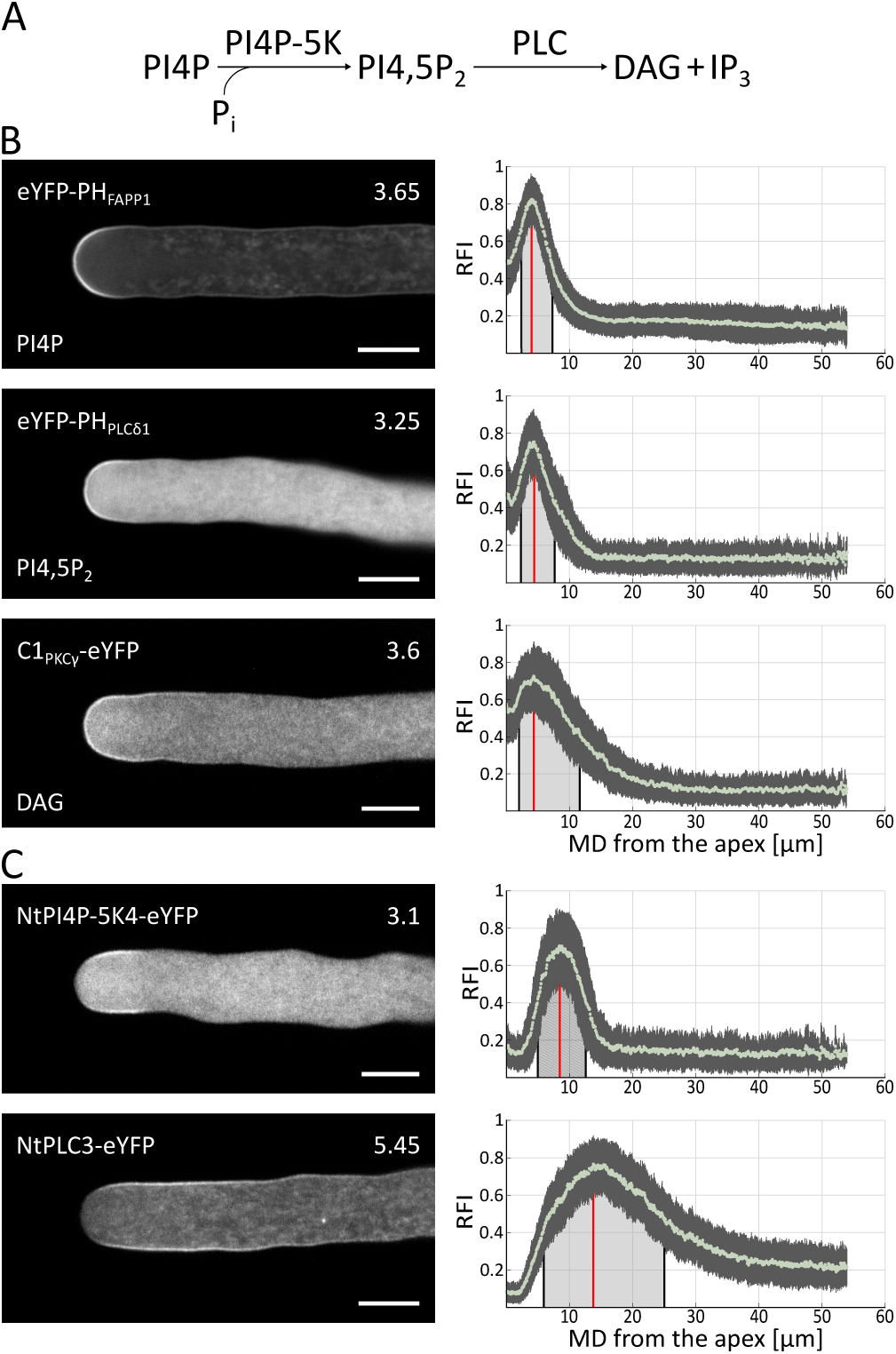
PM domains involved in phosphatidylinositol 4,5-bisphosphate (PI4,5P_2_) metabolism. **A)** Reaction equations representing major pathways of PI4,5P_2_ metabolism. Phosphatidylinositol 4-phosphate 5-kinase (PI4P-5K) activity phosphorylates phosphatidylinositol 4-phosphate (PI4P) to generate PI4,5P_2_. Phospholipase C (PLC) activity hydrolyses PI4,5P_2_ to produce diacylglycerol (DAG) and soluble inositol trisphosphate (IP_3_). **B)**, **C) Left column:** medial confocal optical sections through representative pollen tubes stably expressing the indicated eYFP fusion proteins. Numbers (top right) indicate the growth rate (µm/min) of individual pollen tubes shown, which was determined after image acquisition (average growth rate of all analyzed pollen tubes: Supplemental Fig. S1). Scale bars: 8 µm. **Right column:** quantitative analysis of PM-associated fluorescence (at the PM) in all analyzed pollen tubes expressing each indicated protein. Light green dots: mean relative fluorescence intensity (RFI) associated with the PM at different meridional distances (MDs) from the apex (MD = 0 µm). Both sides of imaged pollen tubes were analyzed, if possible. Dark grey vertical lines: standard deviation. Red vertical lines: maximal RFI. Black vertical lines delimiting light grey shading: proximal and distal half-maximal RFI. **B)** Intracellular distribution and PM association of the indicated eYFP-fusion proteins, which serve as markers for the membrane lipids PI4P (eYFP-PH_FAPP1;_ n = 60 pollen tubes, 116 RFI distribution patterns), PI4,5P_2_ (eYFP-PH_PLCδ1,_ n = 71 pollen tubes, 127 RFI distribution patterns), or DAG (C1_PKCγ1_-eYFP; n = 60 pollen tubes, 120 RFI distribution patterns). **C)** Intracellular distribution and PM association of eYFP fused to the lipid modifying enzymes NtPI4P-5K4 (n = 55 pollen tubes, 104 RFI distribution patterns) or NtPLC3 (n = 58 pollen tubes, 114 RFI distribution patterns).

Phospholipase C (PLC)-mediated PI4,5P_2_ hydrolysis (Fig. 4A) generates the membrane lipid diacylglycerol (DAG) and soluble inositol trisphosphate (IP_3_), both prominent second messengers in animal cells, which activate protein kinase C (PKC) or ER-resident Ca^2+^ channels, respectively (Irvine and Schell, 2001; Kanemaru and Nakamura, 2023). Whether DAG and IP_3_ also have important signaling functions in plants, which do not appear to contain PKC homologs or IP_3_-sensitive Ca^2+^ channels, needs to be elucidated (Meijer and Munnik, 2003). Interestingly, in tobacco and petunia pollen tubes the PLC isoforms NtPLC3 and PetPLC1, respectively, accumulate laterally at the PM in a region flanking the apical PI4,5P_2_ domain and restrict the lateral spreading of this domain (Dowd et al. 2006; Helling et al. 2006).

To gain further insight into the formation and functions of the apical PI4,5P_2_ domain, this domain was quantitatively mapped in normally growing (Supplemental Fig. S1) tobacco pollen tubes alongside PM regions a) enriched in PI4P or DAG (Fig. 4B), or b) associated with enzymes displaying PI4P-5K or PLC activity (Fig. 4C). To visualize PM regions enriched in membrane lipids, the following well-characterized and specific makers were employed, which comprise eYFP fused to different lipid binding domains: a) the PI4P marker eYFP-PH_FAPP1_ (Dowler et al. 2000; Godi et al. 2004), b) the PI4,5P_2_ marker eYFP-PH_PLCδ1_ (Stauffer et al. 1998), and c) the DAG marker C1_PKCγ_-eYFP (Oancea et al. 1998). Each of these markers was previously reported to associate with variable regions of the PM at the tip of transiently transformed tobacco pollen tubes (Kost et al. 1999; Helling et al. 2006; Ischebeck et al. 2008; Thole et al. 2008; Zhao et al. 2010). The relative positions and extensions of the PM domains labeled by these markers were therefore re-examined by quantitative mapping in stably transformed pollen tubes.

Data presented here establish PI4P, PI4,5P_2_ and DAG accumulation within very similar apical PM domains at the tip of tobacco pollen tubes, although the DAG domain distally extended somewhat further than the two other domains (Figs. 4B and 1B). Interestingly, all three domains displayed a pronounced dip in PM-associated fluorescence (Fig. 4B) within a small region at the tip largely overlapping with the apical secretory domain (Fig. 1), with which RAC/ROP^GTP^, NtGEF12a (Fig. 2B) and the RLK AtPRK1 (Grebnev et al. 2020) are strongly and specifically associated. These observations suggest that the accumulation of freely diffusible membrane lipids within the apical secretory domain, with which thousands of vesicles are thought to fuse every minute (Picton and Steer, 1983; Bove et al. 2008; Ketelaar et al. 2008), may be limited by retrograde drift (Fig. 1) and/or molecular crowding (Ellis, 2001; Löwe et al. 2020). By contrast, strong association of RAC/ROP^GTP^, NtGEF12a and AtPRK1 with the apical secretory domain may rely on physical interactions of these proteins with densely packed lipid rafts (Fig. 3B), and/or in case of the TM protein AtPRK1 with the apical cell wall (Grebnev et al. 2020). Importantly, the PI4,5P_2_ domain (Fig. 4B) substantially extends beyond the apical RAC/ROP^GTP^ (Fig. 2B) and secretory domains (Fig. 1A). This demonstrates that although PI4,5P_2_ appears to stimulate apical RAC/ROP activation and secretion as discussed above, additional factors are required to promote these processes and to confine them to the apical secretory domain. Consistent with conserved RHO upstream regulation, apical RAC/ROP activation in tobacco pollen tubes appears to require NtGEF12a-mediated nucleotide exchange in addition to the GDF activity of PI4,5P_2_ (Fig. 2B). Similarly, PI4,5P_2_ may act together with other factors possibly including components of the exocyst (Žarský et al. 2009; Synek et al. 2014) and/or SNARE (Liu et al. 2023) complexes to determine the site of apical secretion.

As discussed above, NtPLC3 (Helling et al. 2006) and PetPLC1 (Dowd et al. 2006) display lateral PM association in growing pollen tubes, although they hydrolyze PI4,5P_2_ to DAG, both of which accumulate at the apex (Fig. 4B; Helling et al. 2006; Scholz et al. 2022). Based on these observations, DAG was suggested to be generated within a lateral PM region, in which the NtPLC3 and PI4,5P_2_ domains overlap (Helling et al. 2006). As DAG presumably is unable to diffuse towards the tip against the retrograde drift within the apical PM (Fig. 1), this lipid was proposed to be internalized within the lateral endocytic domain and to be recycled back to the PM via apical secretion (Helling et al. 2006). Supporting this hypothesis, DAG was observed to accumulate within aberrant endosomal compartments formed in tobacco pollen tubes expressing the phosphatidylinositol 3-phosphate (PI3P) marker RFP-FYVE at high levels (Helling et al. 2006).

Remarkably, different PI4P-5K isoforms were also reported to laterally associate with the pollen tube PM (Ischebeck et al. 2008; Sousa et al. 2008), although both the substrate (PI4P) and the product (PI4,5P_2_) of these enzymes accumulate within apical PM domains in tobacco pollen tubes (Fig. 4B). PI4,5P_2_ therefore appears to be laterally generated within the pollen tube PM where the PI4P-5K and PI4P domains overlap, and similar to DAG may be redistributed to the PM at the tip based on endocytic recycling and apical secretion.

Quantitative analysis of the lateral PM domains associated with eYFP fused to the tobacco pollen tube PI4P-5K isoform NtPI4P-5K4, or to NtPLC3, demonstrated that the proximal ends of both domains were positioned at a similar MD from the apex, although the NtPLC3 domain distally extended much further into the shank (Fig. 4C). Importantly, substantial overlap at the proximal ends of the NtPI4P-5K4 and NtPLC3 domains with distal regions of the apical PI4P and PI4,5P_2_ domains supports direct contact of both enzymes with their substrate within lateral PM regions, which intersect with approximately half of the endocytic domain (Figs. 4B, C and 1B). To further investigate whether PI4,5P_2_ and DAG, which appear to be generated within these lateral PM regions, are in fact endocytically recycled, the distribution of these lipids was imaged in pollen tubes treated with brefeldin A (BFA; Fig. 5A). This drug does not affect endocytosis, but inhibits pollen tube tip growth by blocking the formation of secretory vesicles at the TGN and induces the formation of a single subapical BFA compartment largely comprised of aberrant TGN elements (Nebenführ et al. 2002; Grebnev et al. 2020). The BFA compartment traps PM proteins and lipids, which are delivered to the PM by apical secretion during normal tip growth, either after biosynthesis within the endomembrane system or following endocytic recycling (Geldner et al. 2001; Sheung et al. 2007; Stephan et al. 2014; Grebnev et al. 2020). This can be observed using the styryl dye FM4-64, a commonly employed tracer of endocytic membrane traffic (Grebnev et al. 2020). FM4-64 labels the PM immediately after application, is subsequently endocytically recycled to apical vesicles in normally growing pollen tubes (Fig. 5, DMSO; Parton et al. 2001), and accumulates in the BFA compartment after drug treatment (Fig. 5, BFA; Parton et al. 2001).

**Figure 5:**
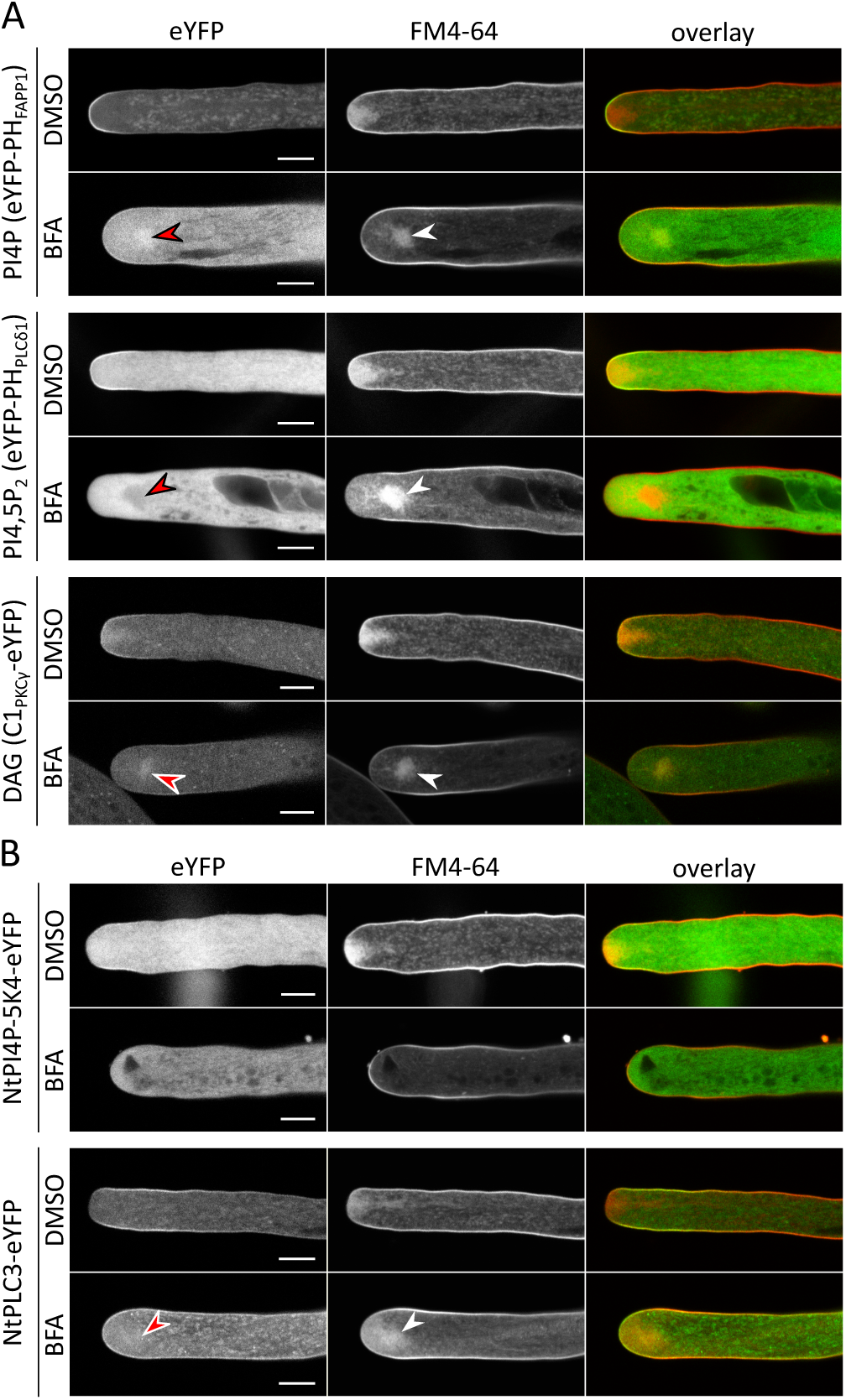
Effects of BFA treatment on the intracellular distribution of lipids and lipid modifying enzymes involved in PI4,5P_2_ metabolism. Medial confocal optical sections through representative pollen tubes stably expressing the indicated eYFP fusion proteins. Pollen tubes were grown in the presence of the lipid dye FM4-64 for 30 min, before BFA was applied for another 20 min. BFA-free solvent was added to control pollen tubes (DMSO). Images showing eYFP (green) or FM4-64 (red) fluorescence were simultaneously recorded in separate channels and overlaid. Control pollen tubes (DMSO) elongated normally at a rate of at least 3 µm/min. BFA treatment stopped pollen tube growth and induced the formation of a subapical BFA compartment, in which apically secreted and endocytically recycled membrane material including FM4-64 labeled lipids (white arrowheads) was trapped. In two independent experiments, all imaged pollen tubes expressing each marker and treated either with DMSO or with BFA showed essentially the same labeling pattern. Number of BFA treated pollen tubes imaged: n = 15 (eYFP- PH_FAPP1_); n = 10 (eYFP-PH_PLCδ1_); n = 11 (C1_PKCγ1_-eYFP); n = 16 (NtPI4P-5K4-eYFP); n = 19 (NtPLC3-eYFP). Scale bars: 8 µm. **A)** In the presence of BFA, eYFP fusion proteins serving as PI4P or DAG markers accumulated in the FM4-64 labeled BFA compartment and remained detectable at the PM, whereas association of the PI4,5P_2_ marker with either of these structures was not detectable (red arrowheads: BFA compartment). **B)** BFA completely abolished PM association of NtPI4P-5K4-eYFP, but not of NtPLC3-eYFP. Like other treatments inhibiting tip growth, BFA application caused NtPLC3-eYFP PM labeling to extend into the apical dome. This fusion protein was also weakly detectable in the BFA compartment (red arrowhead).

DAG not only accumulates in aberrant endocytic organelles induced by RFP-FYVE overexpression as discussed above (Helling et al. 2006), but is also trapped within the BFA compartment (Fig. 5A), strongly suggesting endocytic recycling of this lipid during normal tip growth. By contrast, PI4,5P_2_ accumulation in the BFA compartment was not observed (Fig. 5A). Unexpectedly, BFA treatment completely abolished detectable PM association of NtPI4P-5K4 and the PI4,5P_2_ marker, but not of NtPLC3 and the DAG marker (Fig. 5A and B), indicating that PI4,5P_2_ may not be observed within the BFA compartment because this lipid is no longer generated after drug application. Interestingly, RFP-FYVE overexpression in tobacco pollen tubes was also reported to abolish PI4,5P_2_ detection at the PM (Helling et al. 2006). Currently available data therefore fail to unambiguously establish that PI4,5P_2_ is generated within the lateral PM and endocytically recycled as proposed above. Alternatively, this lipid may be produced by PI4P-5K isoforms, which unlike NtPI4P-5K4 and several of its previously investigated homologs accumulate at the apical pollen tube PM in a region overlapping with the RAC/ROP^GTP^, PI4P and PI4,5P_2_ domains. Indeed, PI4P-5K isoforms have occasionally been reported to associate with the PM at the apex of pollen tubes or root hairs (Kusano et al. 2008; Stenzel et al. 2008, 2020).

Unlike PI4,5P_2_ and DAG, PI4P does not seem to be generated by PM-associated biosynthetic enzymes. PI4P accumulates within the apical PM not only in tobacco pollen tubes (Fig. 4B) but also in Arabidopsis root hairs, in which this lipid is produced by phosphatidylinositol 4-kinase (PI-4K) activity associated with the TGN and with secretory vesicles (Preuss et al. 2006; Thole et al. 2008; Kang et al. 2011). Based on these findings, PI4P accumulating at the pollen tube tip appears likely to also be generated within post-Golgi endomembrane compartments and to be delivered to the PM by apical secretion. Consistent with this hypothesis, PI4P was detectable within the BFA compartment in drug-treated tobacco tubes (Fig. 5A). Whether endocytic recycling also contributed to the observed PI4P accumulation in this compartment remains to be clarified.

Further characterization of the formation and the functions of the apical PI4,5P_2_ domain at the tip of tobacco pollen tubes requires challenging additional investigations. Important research to be pursued includes determining molecular mechanisms responsible for NtPLC3 or NtPI4P-5K4 targeting to specific lateral PM domains, identifying TGN-associated NtPI-4K isoforms, and searching for NtPI4P-5K isoforms that may accumulate at the apical PM.

### PM domains enriched in PA or PS extensively overlap with the lateral NtPLC3 domain

Like PI4,5P_2_, the signaling lipid phosphatidic acid (PA) has various important functions in eukaryotic cells (Testerink and Munnik, 2011; Kim and Wang, 2020; Zhou et al. 2023). PA is generated by diacylglycerol kinases (DGKs) catalyzing DAG phosphorylation, and by phospholipase D (PLD)-mediated hydrolysis of structural membrane lipids. Altering DGK or PLD activity by overexpression, genetic knock-out and/or pharmacological inhibition strongly affects pollen tube tip growth, indicating important roles of PA and of both PA-generating biosynthetic pathways in this process (Pleskot et al. 2012; Vaz Dias et al. 2019; Pejchar et al. 2020; Scholz et al. 2022). Although PA was demonstrated to accumulate in a lateral domain of the tobacco pollen tube PM (Potocký et al. 2014) using an eYFP-tagged highly specific PA-binding domain (eYFP-Spo20p; Nakanishi et al. 2004), the signaling functions of this lipid during tip growth are not understood to date.

As discussed above, PS could potentially promote RAC/ROP^GTP^ nanoclustering (Platre et al. 2019) at the apical PM, which may play an important role in the control of pollen tube tip growth. However, when stably expressed in transgenic Arabidopsis pollen tubes eYFP fused to a specific PS-binding domain (eYFP-C2_Lact_; Yeung et al. 2008) only weakly associated with the apical PM and much more strongly labeled the VAR, in which different RAB family small GTPases displaying specific PS binding *in vitro* also were observed to accumulate (Zhou et al. 2020). Based on these findings, PS was proposed to modulate RAB-controlled apical vesicle trafficking required for Arabidopsis pollen tube growth (Zhou et al. 2020). Supplemental data presented in a report by Platre et al. (2018) also indicated strong eYFP-C2_Lact_ association with apical vesicles in Arabidopsis root hairs and in transiently transformed tobacco pollen tubes. Interestingly, tobacco pollen tubes in addition displayed strong PM labeling along the entire shank, but not at the tip.

Quantitative analysis of PA and PS distribution in normally growing (Supplemental Fig. S1) tobacco pollen tubes stably expressing eYFP-Spo20p or eYFP-C2_Lact_, respectively, established accumulation of both these lipids in lateral PM domains (Fig. 6A), which largely overlap with each other and with the NtPLC3 domain (Figs. 4C and 1B). These observations suggest possible functions of PA and/or PS in NtPLC3 targeting to the lateral PM. Consistent with earlier reports (Platre et al. 2018; Zhou et al. 2020), in tobacco pollen tubes PS also accumulated in cytoplasmic endomembrane compartments including apical vesicles (Fig. 6A), and even in pollen tubes with identical genetic background displayed variable relative levels of accumulation in the lateral PM, in apical vesicles and in other cytoplasmic endomembrane compartments (Supplemental Fig. S5). In rare cases, PS was barely detectable in the PM and strongly accumulated in unidentified cytoplasmic organelles (Supplemental Fig. S5; right image). Importantly, PS was not observed to accumulate in the apical PM of any of the analyzed tobacco pollen tubes. The striking lack of overlap between the lateral PS domain (Figs. 6A and 1B) and the apical RAC/ROP^GTP^ domain (Figs. 2B and 1A) evidently does not support a role of PS in promoting RAC/ROP^GTP^ nanoclustering at the tip of tobacco pollen tubes.

**Figure 6:**
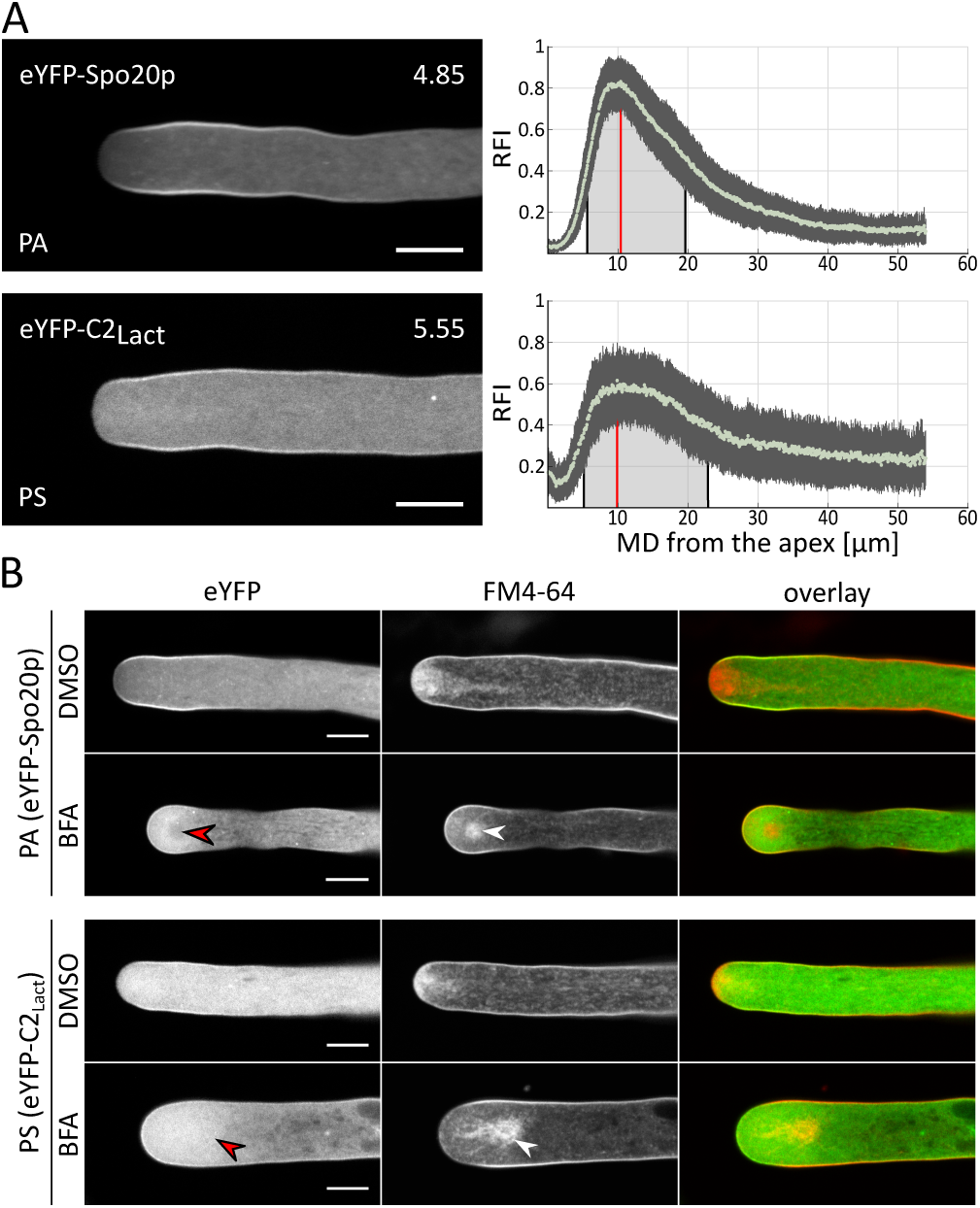
PA- or PS-enriched PM domains and effects of BFA treatment on the intracellular distribution of these lipids. **A) Left column:** medial confocal optical sections through representative pollen tubes stably expressing either the PA marker eYFP-Spo20p or the PS marker eYFP-C2_Lact_. Numbers (top right) indicate the growth rate (µm/min) of individual pollen tubes shown, which was determined after image acquisition (average growth rate of all analyzed pollen tubes: Supplemental Fig. S1). Scale bars: 8 µm. **Right column:** quantitative analysis of PM-associated fluorescence (at the PM) in all analyzed pollen tubes expressing eYFP-Spo20p (n = 58 pollen tubes, 115 RFI distribution patterns) or eYFP-C2_Lact_ (n = 60 pollen tubes, 115 RFI distribution patterns). Both sides of imaged pollen tubes were analyzed, if possible. Light green dots: mean relative fluorescence intensity (RFI) associated with the PM at different meridional distances (MDs) from the apex (MD = 0 µm). Dark grey vertical lines: standard deviation. Red vertical lines: maximal RFI. Black vertical lines delimiting light grey shading: proximal and distal half-maximal RFI. **B)** Medial confocal optical sections through representative pollen tubes stably expressing the indicated PA or PS marker. Pollen tubes were grown in the presence of the lipid dye FM4-64 for 30 min, before BFA was applied for another 20 min. BFA-free solvent was added to control pollen tubes (DMSO). Images showing eYFP (green) or FM4- 64 (red) fluorescence were simultaneously recorded in separate channels and overlaid. Control pollen tubes (DMSO) were normally elongating at a rate of at least 3 µm/min. BFA treatment stopped pollen tube growth and induced the formation of a subapical BFA compartment, in which apically secreted and endocytically recycled membrane material including FM4-64 labeled lipids (white arrowheads) was trapped. Whereas PA clearly was excluded from the FM4-64 labeled BFA compartment, PS accumulated in this compartment, albeit not to levels beyond the bright cytoplasmic background (red arrowheads). In two independent experiments, all imaged pollen tubes expressing each marker and treated either with DMSO or with BFA showed essentially the same labeling pattern. Number of BFA treated pollen tubes imaged: n = 12 (eYFP-Spo20p); n = 8 (eYFP-C2_Lact_). Scale bars: 8 µm.

PS biosynthesis commonly occurs within the endomembrane system of both animal and plant cells (Vance and Steenbergen, 2005; Yamaoka et al. 2011), suggesting that this lipid is delivered to the pollen tube PM by apical secretion. Consistent with this assumption, PS was detected in the BFA compartment of drug-treated tobacco pollen tubes, although it did not accumulate in this compartment to levels beyond the bright cytoplasmic background (Fig. 6B), which appears to represent labeling of endomembrane compartments. Considerable overlap between the lateral PS and endocytic domains (Figs. 6A and 1B) indicates that endocytic recycling may contribute to PS accumulation in the BFA compartment. However, as in the case of PI4P, which also appears to be generated within the endomembrane system and apically secreted, endocytic recycling of PS needs to be confirmed using a different approach.

Unlike PS, PA appears to be generated laterally at the pollen tube PM, where this lipid accumulates (Fig. 6A; Potocký et al. 2014) together with different tobacco pollen tube DGK (Scholz et al. 2022) and PLD (Pejchar et al. 2020) isoforms. Remarkably, PA was not detectable in the BFA compartment after drug treatment (Fig. 6B), although this lipid a) massively accumulates within the lateral endocytic domain in normally growing pollen tubes (Figs. 6A and 1B), and b) by contrast to PI4,5P_2_ remained detectable at the PM after BFA application. Therefore, PA appears to be excluded from lateral endocytic recycling, strongly suggesting that not only TM proteins (Grebnev et al. 2020) but also membrane lipids are selectively internalized during this process based on specific recognition mechanisms. By contrast, pollen tube PM lipids were previously proposed to undergo constitutive endocytic recycling (Grebnev et al. 2020).

To further characterize PA and PS functions in the control of tip growth, it will be necessary to identify pollen tube proteins that interact with these lipids and to determine whether NtPLC3 is among them. If so, it may be possible to establish essential functions of PS and/or PA in NtPLC3 targeting to the lateral PM e.g. by manipulating the activity of PLDs (Potocký et al. 2003, 2014; Pejchar et al. 2020;), DGKs (Scholz et al. 2022), or other enzymes required for the biosynthesis of these lipids in tobacco pollen tubes. Characterization of cellular and molecular mechanism responsible for PS and PA targeting to the lateral pollen tube PM will improve our understanding of the regulation of tip growth by these lipids. A first step towards this goal will be the quantitative mapping of PM domains associated with endogenous DGK and PLD isoforms in normally growing tobacco pollen tubes using the methodology introduced here.

### Integrated analysis of PM and cytoplasmic compartmentalization indicates subapical diffusion barriers and further defines Ca^2+^ functions in apical secretion

The mean MDs from the pollen tube apex of the proximal (equals 0 for all apical PM regions) and distal endpoints of all protein and lipid PM domains investigated here were determined and compared based on statistical analysis (Fig. 7A). Figure 7A shows a linear alignment of all PM domains quantitatively characterized in this manner. For direct comparison, the apical PM domain displaying high membrane order, which is identical to the apical secretory domain (Fig. 1), as well as the local peak in membrane order flanking this domain, are also indicated in this figure, although the endpoints of these two PM regions were either predefined or distinctly determined as described above (Fig. 3). Remarkably, the alignment reveals that the distal ends of the apical RAC/ROP^GTP^ and NtGEF12a domains, as well as the proximal ends of all lateral protein (NtGAP1, NtPI4P-5K4, NtPLC3) and lipid (PA, PS) domains, are positioned within a narrow PM region at the base of the apical domain, which extends from ca. 3.0 - 6.3 µm MD from the apex (Fig. 7A; light blue shading). Remarkably, this region proximally coincides with the local peak in membrane order (Figs. 7A and 3C - E), and extensively overlaps with the previously reported (Grebnev et al. 2020) PM contact domains of the F-actin fringe and of the TGN compartment, which are located in the subapical cytoplasm (Fig. 1). To confirm this important finding, the PM contact domains of these two cytoplasmic structures were reassessed based on the procedures employed here to map protein and lipid domains (Fig. 7B).

**Figure 7:**
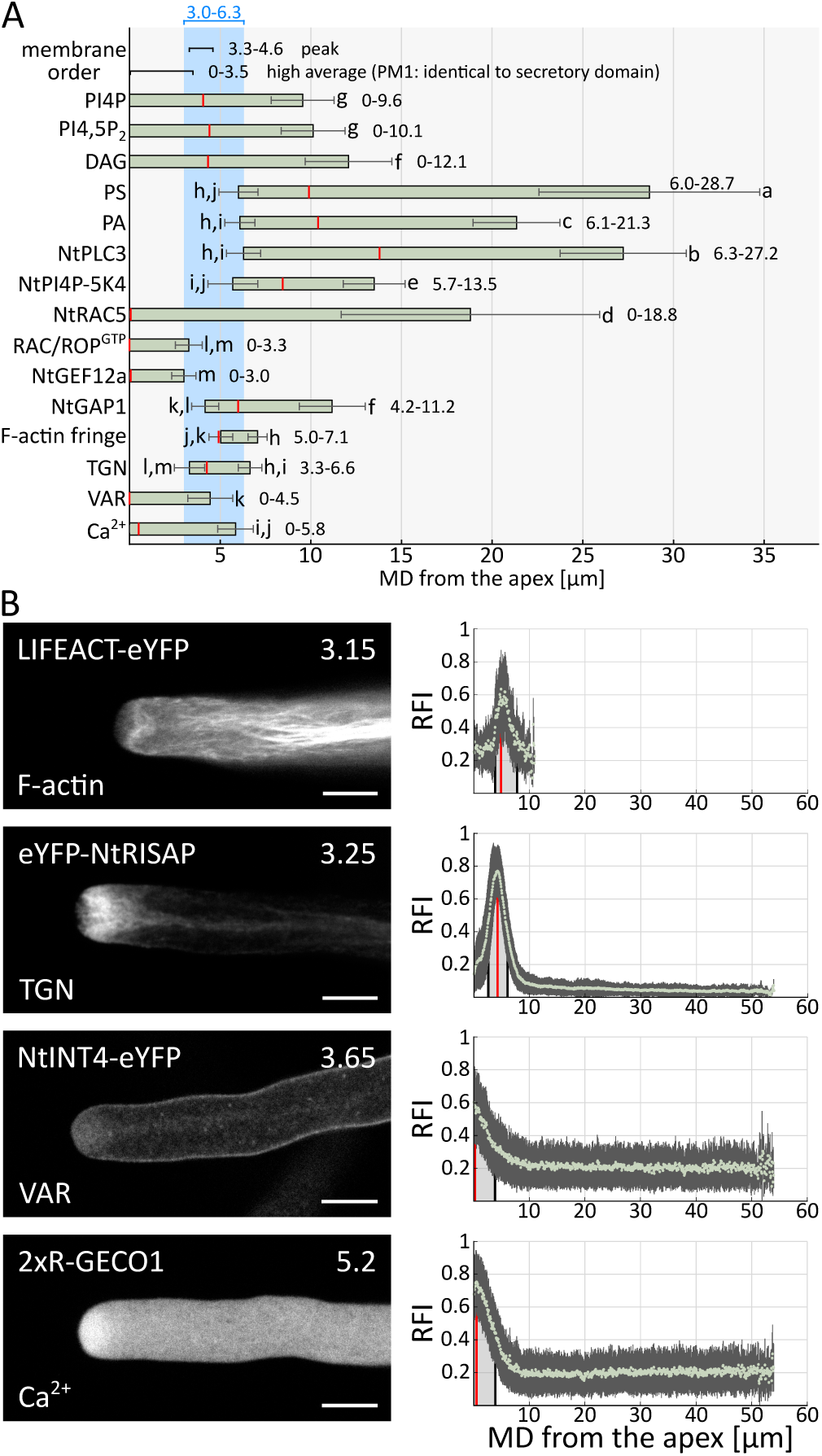
Comparative quantitative analysis of PM and cytoplasmic compartmentalization. **A)** Alignment of linear bars representing all PM domains associated with various proteins and lipids (Figs. 2, 4 and 6A), or in contact with different cytoplasmic structures or regions (Fig. 7B), which were investigated in this study. Mean meridional distances (MDs) from the pollen tube apex (MD = 0 µm) of the proximal (if applicable) and distal endpoints of these PM domains were quantitatively determined (values indicated next to each bar and in Fig. 1) and statistically compared using one-way ANOVA with Tukey Kramer (HSD) testing. Error bars: standard deviation. Distinct letters indicate statistically significantly different mean MDs from the apex (p ≤ 0.05). Red vertical lines: maximal RFI indicated in figures 2, 4, 6A and 7B. Number of pollen tubes/domain endpoints analyzed (both sides of pollen tubes were analyzed, if possible): PI4P, n = 70/140; PI4,5P_2_, n = 83/155; DAG, n = 65/130; PS, n = 60/119; PA, n = 81/162; NtPLC3, n = 117/231; NtPI4P-5K4, n = 55/106; NtRAC5, n = 55/107; RAC/ROP^GTP^, n = 66/132; NtGEF12a, n = 61/122; NtGAP1, n = 67/134; F-actin fringe, n = 63/126; TGN, n = 69/138; VAR, n = 57/114; Ca^2+^, n = 67/134. For direct comparison, the extension of the apical PM domain displaying high membrane order as well as the position of the lateral peak in membrane order are also indicated (Fig. 3). The apical PM domain displaying high membrane order is identical to the secretory domain. Light blue shading indicates a PM region in which the endpoints of many apical and lateral domains are positioned. All domains are drawn to scale. **B) Left column:** medial confocal optical sections through representative pollen tubes stably expressing the indicated fluorescent fusion proteins serving as markers for the following cytoplasmic structures or regions: F-actin (LIFEACT-eYFP), subapical TGN compartment (eYFP-RISAP), VAR (NtINT4-eYFP) or region displaying elevated Ca^2+^ concentrations (2xR-GECO1). Numbers (top right) indicate the growth rate (µm/min) of individual pollen tubes shown, which was determined after image acquisition (average growth rate of all analyzed pollen tubes: Supplemental Fig. S1). Scale bars: 8 µm. **Right column:** quantitative analysis of PM-associated fluorescence (directly underneath the PM) in all analyzed pollen tubes expressing LIFEACT-eYFP (n = 59 pollen tubes, 110 RFI distribution patterns), eYFP-NtRISAP (n = 58 pollen tubes, 116 RFI distribution patterns), NtINT4-eYFP (n = 55 pollen tubes, 109 RFI distribution patterns) or 2xR-GECO1 (n = 55 pollen tubes, 108 RFI distribution patterns). Both sides of imaged pollen tubes were analyzed, if possible. Light green dots: mean relative fluorescence intensity (RFI) associated with the PM at different meridional distances (MD) from the apex (MD = 0 µm). Dark grey vertical lines: standard deviation. Red vertical lines: maximal RFI. Black vertical lines delimiting light grey shading: proximal (if applicable) and distal half-maximal RFI.

Previously reported data (Fig. 1; Grebnev et al. 2020) and results presented here (Fig. 7) identified similar endpoints of the PM region in contact with the subapical TGN, which was labeled by the eYFP-tagged myosin receptor NtRISAP (Stephan et al. 2014). By contrast, both endpoints of the PM domain associated with the LIFEACT-eYFP-labeled F-actin fringe (Riedl et al. 2008; Montes-Rodriguez and Kost, 2017) determined here (Fig. 7) appeared to be shifted 1.4 - 1.5 µm in distal direction when compared to previously reported data (Fig. 1A; Grebnev et al. 2020). As this discrepancy cannot be attributed to differences in the average growth rate of the analyzed pollen tubes, which was the same in both studies (Supplemental Fig. S1; Grebnev et al. 2020), it appears to be a consequence of the application of distinct procedures to analyze PM association of the notoriously weakly labeled and rapidly bleaching F-actin fringe. In any case, taken together the two currently available data sets indicate association of the F-actin fringe with a PM region extending from 3.6 - 7.1 µm MD from the apex (Fig. 1A). This region extensively overlaps with the PM domain, within which the distal or proximal endpoints of different apical and lateral domains, respectively, are located, as discussed in the previous paragraph (Fig. 7A; light blue shading).

In different cell types, cortical F-actin structures interact with peripheral or integral membrane proteins, and/or with membrane lipids (lipid pinning; Fujimoto and Parmryd 2017), to specifically restrict the lateral mobility of these PM components and of their interaction partners (Andrade et al. 2015; Jaqaman and Grinstein 2012; Nakada et al. 2003). Furthermore, tight association of cortical actin filaments with the PM results in space restrictions creating a physical barrier, which further limits lateral mobility within the PM (Nakada et al. 2003). Interestingly, lipid pinning by PM-associated actin filaments can also induce the formation of highly ordered PM domains (Dinic et al. 2013). In the light of these findings, it is conceivable that the pollen tube F-actin fringe establishes a local diffusion barrier at the base of the apical dome, which supports the separation between apical and lateral PM domains. Actin-induced formation of the local peak in membrane order within the PM, which overlaps with the proximal end of the F-actin fringe (Figs. 1, 3D and E), may contribute to this diffusion barrier. However, signaling lipids with relatively small cytoplasmic head groups, which are delivered to the PM by apical secretion, generally appear to be able to pass this diffusion barrier (Fig. 7A) and to either form apical PM domains that distally extend beyond the F-actin fringe (PI4P, DAG and possibly PI4,5P_2_), or to accumulate in a lateral PM domain (PS). Whereas spreading of the apical PI4P and PI4,5P_2_ domains appears to be limited by PI4P-5K and PLC isoforms associated with the lateral PM, respectively, retrograde transport of PS within the PM does not seem to be similarly confined by lipid modifying enzymes.

The subapical TGN compartment, which is held in place by the F-actin fringe and integrates apical membrane traffic as outlined above (Stephan et al. 2014), appears to extend the proposed diffusion barrier within the PM into the cytoplasm. This TGN compartment was demonstrated to separate the apical VAR, which is densely packed with secretory vesicles, from the cytoplasm in the pollen tube shank that contains all other cell organelles (Stephan et al. 2014). In tobacco pollen tubes, these two cytoplasmic regions also display striking differences in F-actin organization (Fig. 7B): whereas the VAR contains little or no F-actin (Montes-Rodriguez and Kost, 2017), a dense network of largely longitudinally oriented F-actin cables drives massive cytoplasmic streaming in the pollen tube shank, along which most organelles are rapidly moving (Cai et al. 2015).

To further characterize the confinement of the VAR by the subapical TGN compartment, the PM contact domain of the VAR was quantitatively determined based on labeling apical vesicles with eYFP fused to NtINT4 (Fig. 7A and B), an endogenous tobacco pollen tube TM protein delivered to the PM by apical secretion (Grebnev et al. 2020). Interestingly, the apical PM domain associated with the VAR distally extended somewhat further than the secretory domain and overlapped to some extent with the proximal end of the TGN domain (Figs. 1B and 7A). This indicates that apical vesicles may intermingle with TGN elements at the edge of the VAR, where they seem to be unable to fuse with the PM. Consistent with previous reports (Parton et al. 2001; Cai et al. 2015), the subapical TGN compartment did not completely prevent loss of apical vesicles from the VAR. In the cell center at the distal tip of the VAR, which deeply penetrates into the torus-shaped subapical TGN compartment (Fig. 1; Stephan, 2017), NtINT4-eYFP-containing apical vesicles were observed to escape from the VAR into the cytoplasm of the shank, where they joined retrograde cytoplasmic streaming (Fig. 7B). The possible functional relevance of this phenomenon remains to be determined.

Extensive evidence supports an essential but only partially understood role of a tip-focused Ca^2+^ gradient in the control of apical secretion, which drives the polar expansion of pollen tubes and other types of tip-growing cells (Robert Konrad et al. 2018; Tian et al. 2020). Stable expression of the intensiometric Ca^2+^ sensor 2xR-GECO1 (Zhao et al. 2011; Li et al. 2021) enabled detection of a cytoplasmic region at the tip of normally growing tobacco pollen tubes (Supplemental Fig. S1), which exhibited elevated Ca^2+^ concentrations (Fig. 7B). In individual growing pollen tubes, the extension of this region, as well as 2xR-GECO1 fluorescence intensity within its borders, remained constant over time. Consistent with this observation, neither the growth rate nor the steepness of the tip-focused Ca^2+^ gradient are necessarily oscillating during normal pollen tube tip growth under optimal conditions (Chebli and Geitmann, 2007; Iwano et al. 2009; Hemelryck et al. 2017).

Interestingly, the apical cytoplasmic region exhibiting elevated Ca^2+^ concentrations largely overlapped with the NtINT4-eYFP-labeled VAR (Fig. 7B). However, quantitative analysis revealed that the PM domain in contact with the cytoplasmic region displaying elevated Ca^2+^ concentrations distally extended a little further than the VAR-associated PM domain (Fig. 7A), which in turn was somewhat larger than the apical secretory domain, as discussed in the previous paragraph (Fig. 1B). Together, these findings suggest that Ca^2+^ does not determine the lateral extension of the apical secretory domain in tobacco pollen tubes by directly triggering vesicle fusion with the apical PM. In contrast, elevated Ca^2+^ concentrations may prime secretory vesicles throughout the VAR for fusion with the PM, similar to what has been observed in mammalian neuronal cells (Martin, 2012; Ammar et al. 2013; Frere et al. 2015).

Further investigation is clearly required of the proposed diffusion barrier at the base of the apical dome, the dynamic behavior of apical vesicles within the VAR and the fusion of these vesicles with the PM. Proteins associated with the F-actin fringe and with the lateral peak in membrane order need to be identified and functionally characterized. In addition, the mobility of proteins and lipids accumulating in different PM domains, or in apical vesicles, needs to be quantitatively analyzed based on FCS, single molecule tracking and labeling with photoactivatable (or photoconvertible) fluorescent proteins.

### Conclusions

Data presented here quantitatively define an elaborate dynamic structural framework, which is essential for tobacco pollen tube tip growth. Remarkable aspects of this framework provide important insights into regulatory and cellular processes required for directional cell expansion, and delineate important areas of further research:

- NtGEF12a-mediated RAC/ROP activation appears to determine the site of apical secretion.
- Membrane order is high within the apical secretory PM domain and potentially promotes RAC/ROP^GTP^ nanoclustering essential for downstream signaling.
- PI4,5P_2_, which stimulates RAC/ROP PM association and can promote vesicle fusion with the PM, exerts these functions together with other factors, which determine exact sites of RAC/ROP activation and apical secretion.
- Key signaling lipids generated either within the endomembrane system or at the PM accumulate in distinct PM domains depending on membrane trafficking, retrograde drift within the PM and the activity of PM-associated lipid modifying enzymes.
- Like TM proteins, membrane lipids appear to be selectively internalized by lateral endocytosis.
- PA and PS extensively colocalize with NtPLC3, suggesting important functions of these lipids in specific NtPLC3 targeting to the lateral PM.
- PS does not overlap with RAC/ROP^GTP^ and therefore cannot promote potential nanoclustering of this protein.
- The F-actin fringe, a local membrane order peak within the PM and the subapical TGN compartment appear to form a diffusion barrier at the base of the apical dome, which promotes the separation of apical and lateral PM domains, and restricts exchange between the apical VAR and the rest of the cytoplasm.
- Ca^2+^ levels are elevated throughout the apical VAR and do not appear to directly trigger vesicle fusion with the PM, but may prime secretory vesicles for this process.

## Material and methods

### Plasmids

Standard techniques (Sambrook et al. 1989) were employed for plasmid construction. To generate plant expression vectors, PCR-amplified cDNA sequences were inserted into the multiple cloning site (MCS) of pHD32 (containing a p*Lat52*:MCS-5xGA-*eYFP*-*Nos*T cassette) or of pWEN240 (containing a p*Lat52*:*eYFP*-5xGA-MCS-*Nos*T cassette) (Grebnev et al. 2020; Klahre et al. 2006). Resulting expression vectors contain cDNA sequences, which encode eYFP fused via a flexible quintuple glycine/alanine linker (5xGA) to the N- or C-terminus of different proteins or protein domains. Expression of these cDNA sequences is controlled by an upstream pollen-specific *Lat52* promotor (p*Lat52*; Twell et al. 1990) and a downstream *Nos* terminator (*Nos*T; Bevan et al. 1983). All plant expression vectors and the expression cassettes they contain are listed in supplemental table S1, along with the PCR templates, PCR primers and restriction enzymes used to construct them. Supplemental table S2 contains further information about plasmids employed as PCR templates. PCR primer sequences are provided in supplemental table S3.

To generate binary vectors enabling *Agrobacterium*-mediated plant transformation, expression vectors listed in supplemental table S1 were restricted to release cDNA sequences encoding eYFP fusion proteins. These cDNA sequences were inserted between the *Lat52* promoter and the *Nos* terminator into the MCS of pHD71 (Fritz and Kost 2020), a derivative of the binary vector pPZP212 (Hajdukiewicz et al. 1994).

Plasmids employed for split-ubiquitin assays were constructed using plant expression vectors listed in supplemental table S1 as PCR templates. To generate pMetOYC-NtGEF12a, the *NtGEF12a* coding sequence was amplified from pFAU422, inserted into *Bam*HI/*Xho*I-restricted pENTR2B (ThermoFischer; A10463) and transferred from there into a pMetOYC destination vector (Karnik et al. 2015) based on Gateway LR recombination. To generate pNubG-NtRAC5, the *NtRAC5* coding sequence was amplified from pHD23, inserted into *Bam*HI/*Xho*I-restricted pENTR3C (ThermoFischer; A10464) and transferred from there into an pNX35-Dest destination vector (Grefen et al. 2007) based on Gateway LR recombination.

### Plant material

Wild-type and transgenic tobacco (*Nicotiana tabacum* Petit Havana SR1) plants were grown in soil (Hawita) under long day conditions (16 h light at 22°C; 8 h darkness at 18 or 22°C) in a growth chamber or in a green house, which were set to a constant relative humidity of 60 %. Seeds were germinated in soil and emerging seedlings were individually transferred to larger pots after 2 - 3 weeks. Pollen was harvested from mature flowers and generally used to establish pollen tube cultures immediately after collection. Alternatively, pollen-containing anthers were transferred into gelatin capsules (https://kapselwelt.de), frozen in liquid nitrogen and stored at - 80°C for later use.

To establish axenic cultures, tobacco seeds were sterilized in a microwave oven and transferred onto solid (0.8 % phyto agar [Duchefa]) half-strength Murashige Skoog (MS) medium (Duchefa) containing 2 % sucrose (Roth). These cultures were kept in a growth chamber under long day conditions (16 h light at 22°C; 8 h darkness at 18°C) and used for *Agrobacterium*-mediated plant transformation as well as for RT-PCR analysis of gene expression in roots of 7 d-old seedlings.

### Stable tobacco transformation

Transgenic tobacco lines were established essentially as previously described (Horsch et al. 1985). Young leaves of 4 - 5 week-old wild-type tobacco plants grown under axenic conditions were cut into pieces, which were incubated for 10 - 15 min at RT submerged in a suspension of *Agrobacterium tumefaciens* AGL1 (Lazo et al. 1991) cells containing binary expression vectors. Leaf pieces were then placed bottom-side-down onto solid (0.8 % phyto agar [Duchefa]) sugar-free full-strength MS medium and incubated for 2 - 3 days at 21°C in the dark. Subsequently, leaf pieces were transferred onto solid (0.8 % phyto agar [Duchefa]) MGI medium (full-strength MS medium containing 1.6 % glucose [Roth], 1 mg/l BA [Sigma-Aldrich], 0.2 mg/l NAA [Sigma-Aldrich], 100 mg/l Kanamycin [Roth] and 500 mg/l Ticarcillin [Duchefa]) and incubated in a growth chamber under long day conditions (16 h light at 22°C; 8 h darkness at 18°C). To promote callus formation, leave pieces were weekly sub-cultured onto fresh MGI medium. After 3 - 4 weeks, callus-forming leaf pieces were transferred onto fresh MGI medium in tall culture containers. To induce root formation, shoots emerging from calli 3 - 4 weeks later were transferred onto solid (0.8 % phyto agar [Duchefa]) full-strength MS medium containing 2 % sucrose and 250 mg/l Ticarcillin. Shoots with well-developed roots were transferred to soil. Pollen produced by primary regenerants was germinated *in vitro* and emerging pollen tubes were screened for eYFP emission using a Leica DMI4000B fluorescence microscope (excitation: 490 - 510 nm; emission: 520 - 550 nm; dichroic: LP 515).

### *In vitro* pollen germination and pollen tube culture

To establish pollen tube cultures for microscopic analysis, anthers collected from 1 - 3 tobacco flowers were suspended in 15 ml liquid PTNT medium (Read et al. 1993a; 1993b) and pollen was released by vigorous vortexing. Frozen anthers stored at −80°C were suspended in PTNT medium at 4°C and kept at this temperature for 10 minutes before vortexing. After removing empty anthers, pollen contained in 5 ml suspension was vacuum-filtrated onto a round polyamide membrane filter (diameter 5 cm, pore size 0.45 µm; Sartorius) and transferred onto 3.5 ml solid (0.25 % phytagel [Sigma-Aldrich]) PTNT medium in a 5.5 cm diameter Petri dish by briefly placing the filter upside-down onto the medium surface, as previously described (Fritz and Kost 2020; Johnson and Kost 2010). After sealing Petri dishes with Parafilm^®^, cultures were incubated at 21°C in the dark for at least 2 - 3 hours before microscopic analysis.

For transient transformation, pollen tube cultures were established as described in the previous paragraph with the follow modification: anthers collected from 2 tobacco flowers were suspended in 5 ml liquid PTNT medium.

### Transient gene expression in pollen tubes

Newly established pollen tube cultures were immediately transiently transformed by ballistic particle bombardment as previously described (Fritz and Kost 2020; Johnson and Kost 2010; Sun et al. 2015). Briefly, a Bio-Rad PDS-10000-He biolistic particle delivery system (Bio-Rad) and 1000-1100 psi rupture disks were employed to bombard pollen tube cultures placed in a vacuum chamber (28 inches Hg) with 1.6 μm gold particles, which had been coated with 3 - 5 µg plasmid DNA in the presence of CaCl_2_ and protamine.

### FM4-64 staining and brefeldin A (BFA) treatment of cultured pollen tubes

Newly established pollen tube cultures were incubated for 2.5 h at 21°C in the dark, before 200 µl PTNT medium containing 50 µM FM4-64 (ThermoFisher; stock solution: 10 mM in DMSO) was added and cultures were incubated for another 30 min at RT. Subsequently, 200 µl PTNT medium containing 70 µM BFA (ThermoFisher; stock solution: 10 mM in DMSO) was added and cultures were incubated for another 20 min at RT. Identically treated pollen tube cultures, to which 200 µl PTNT medium containing no BFA but DMSO diluted to the same concentration, served as solvent controls. Pollen tubes were imaged by confocal microscopy within 60 minutes after the last 20 min incubation period.

### Confocal microscopy

Sectors of pollen tube cultures were cut out, transferred upside-down onto a coverslip and observed under an inverted Leica SP8 DIVE FALCON laser scanning microscope using an HCX PL APO CS 63.0x/1.20 NA water immersion objective. Pinhole size was adjusted to 1 airy unit. A constant zoom factor of 2.5x was employed, resulting in a pixel size of 72 x 72 nm. Photomultiplier gain and transmission filters were set to maximize image brightness, avoiding overexposure by working in the QLUT (Quick Look Up Table) mode provided by the Leica LAS X software. Additional imaging parameters are indicated in supplemental table S4. eYFP and FM4-64 fluorescence was excited using a 514 nm argon laser. eYFP emission was detected within a 525 - 585 nm window during single-channel imaging, and within a 525 - 575 nm window during sequential dual-channel imaging together with FM4-64 emission, which was detected within a 650 - 790 nm window. 2xR-GECO1 fluorescence was excited using a 561-nm laser line and detected within a 570 - 651 nm emission window. To facilitate the identification of medial planes for confocal imaging, transmitted light reference images were simultaneously acquired. To enable determination of the growth rate of imaged pollen tubes after the acquisition of confocal sections employed for image analysis, a second confocal section was recorded two minutes later.

### Determination of pollen tube growth rates, PM-associated relative fluorescence intensities (RFIs) and PM domain endpoints

The Fiji version of ImageJ (Schindelin et al. 2012) was employed for all these analyses. To measure the growth rates of individual pollen tubes after the acquisition of fluorescence images employed for data analysis, acquired images were overlayed with a second image of the same pollen tube recorded 2 minutes later using identical microscope settings (Fritz and Kost 2020; Johnson and Kost 2010; Sun et al. 2015). The distance in μm between the extreme pollen tube apices on the overlaid images was determined and divided by 2, yielding pollen tube growth rates in μm/min. Pollen tubes with growth rates lower than 3.0 µm/min were excluded from further data analysis. Box plots displaying average growth rates of all analyzed pollen tubes were generated using an R-based (Spitzer et al. 2014) online tool (http://shiny.chemgrid.org/boxplotr/).

Before determination of PM-associated RFIs or PM domain endpoints, all images were converted to 8 bit dynamic range, and “rolling ball” background subtraction was generally performed using a radius of 18 or 50 pixel, respectively. In individual cases PM-associated RFIs were determined without background subtraction (free eYFP) or using “rolling ball” background subtraction with a radius of 50 pixel (2xR-GECO1).

To determine PM-associated RFIs, the “segmented line” tool was employed to draw a 1 pixel-wide line onto background-subtracted images starting at the extreme apex of analyzed pollen tubes and following their perimeter towards the shank. This line delineated either a) the PM associated with eYFP-tagged peripheral membrane proteins or markers for membrane lipids, or b) the cortical cytoplasm directly underneath the PM, which was labeled by eYFP, or by markers for cytoplasmic regions or structures. Using the “profile” tool, fluorescence intensities displayed by each pixel along the segmented line were exported into Excel (Microsoft), and pixel positions were converted to meridional distances (MDs) from the apex (MD = 0) in μm. Resulting intensity profiles were normalized and turned into RFI profiles by assigning the value 1 to the highest fluorescence intensity (Chebli et al. 2012). RFI profiles obtained from all analyzed pollen tubes were averaged to generate plots displaying mean PM-associated RFIs at different MDs from the apex, which are shown in figures 2, 4, 6A and 7B. Maximal and half-maximal mean PM-associated RFIs are listed in supplemental table S5 and are also indicated in these plots.

The “segmented line” tool was also employed to draw lines along the perimeter of analyzed pollen tubes, which connected the extreme apex to the proximal or the distal endpoints of analyzed PM domains. To facilitate the identification of these endpoints, the 16 color Look-Up Table (LUT) was applied. The length of the segmented lines in μm corresponded to the MD from the apex of domain endpoints. Mean MDs from the apex of the endpoints of all analyzed PM domains are listed in supplemental table S6, and are indicated in figures 1 and 7A. Standard deviations are provided in supplemental table S6.

### Di-4-ANEPPDHQ staining and imaging by widefield microscopy

To assess membrane order, pollen tubes were stained with Di-4-ANEPPDHQ, which specifically labels the outer leaflet of the plasma membrane (Dinic et al. 2013). Frozen pollen grains collected from two anthers were rehydrated in 20 ml liquid PTNT medium for 15 min at 4°C, before 1.5 ml of pollen suspension was transferred onto a filter membrane and from there onto 1.8 ml PTNT medium solidified with 0.25 % phytagel (Sigma) in a 5.5 cm Petri dish essentially as previously described (Kost et al. 1998). Plates were incubated at RT in the dark. After 3 h, 200 µl liquid PTNT medium containing 50 µM Di-4-ANEPPDHQ (ThermoFisher; stock solution 10 mM in EtOH) were added and plates were incubated for another 15 min at RT in the dark. Subsequently, squares of solid culture medium covered with Di-4-ANEPPDHQ-stained pollen tubes were cut out and flipped upside down onto a coverslip. Widefield fluorescence imaging was performed using a custom-built system (Supplemental Fig. S6) based on a Nikon Ti-U inverted microscope (Nikon) equipped with a 60x water objective (NA 1.27), a FF01-387/480 dual bandpass excitation filter (Semrock) and a zt405/473-491/NIRrpc dichroic mirror (Chroma). Di-4-ANEPPDHQ was excited at 470 nm using a pE-2 LED light source (CoolLED). Fluorescence emission was split into two channels (510 - 550 nm and ≥ 645 nm) using a dual viewer (Photometrics) equipped with a 585DCXR dichroic mirror (Chroma), as well as with 61003m (510 - 550 nm) and HQ645lp (≥ 645 nm) emission filters (Chroma). Images were acquired using a Cascade 1k CCD camera (Photometrics). To minimize dye photobleaching, focus adjustment was performed without Di-4-ANEPPDHQ excitation using transmitted light with UV blocked by a GG455 long pass filter (Chroma). Stacks of eleven images with a Z-step size of 200 nm and 91 nm square pixels were acquired around the medial pollen tube plane in both emission channels simultaneously. After 2 minutes, a second set of Z-stacks was recorded to enable growth rate determination.

### Deconvolution of Di-4-ANEPPDHQ image stacks

Image stacks acquired in both channels as described in the previous section were simultaneously deconvolved using Huygens Professional (SVI) to improve resolution prior to generalized polarization (GP)-analysis (Huang et al. 2023). Signal to noise (SN) ratio settings ranging from 1 to 5 were used to optimize results. The number of iterations was set to a maximum of 50 and deconvolved images acquired in each of the two channels were saved using a linked scale, which maintains the relative intensity between the two channels. Based on visual inspection of deconvolved images acquired in the noisier lower channel (510 - 550 nm), images displaying the highest SN ratio without signs of over-restoration were selected for subsequent GP-analysis together with corresponding deconvolved images recorded in the upper (≥ 645 nm) channel.

### Determination of general polarization (GP)

Deconvolved Di-4-ANEPPDHQ images generated as described in the previous two sections, and the Fiji version of ImageJ (Schindelin et al. 2012) enhanced with specifically designed macros, were employed for GP-analysis and to determine the growth rate of pollen tubes after the acquisition of images used for this analysis. To semi-automatically delineate the pollen tube PM, a few sequential points along the PM were roughly marked and, assuming that fluorescence intensity peaks at the PM, accurately repositioned by searching for the highest-intensity pixel along a short line segment positioned perpendicularly to the local orientation of the PM. Additional points were interpolated and similarly optimized. The delineated PM was used to automatically identify the pollen tube apex. An area outside each analyzed pollen tube was automatically defined and used to measure the background in each channel, which was subtracted. Five different biologically relevant regions of interest (ROIs) were selected for automatized GP-analysis. The PM at the tip was divided into three one-pixel-wide ROIs spanning distinct MD (indicated in brackets) from the apex: PM1 (0 - 3.5 µm), PM2 (3.5 - 8 µm), and PM3 (8 - 15 µm). Two additional ROIs were defined covering cytoplasmic regions three pixels below the PM underneath either PM1 or PM3: 1) the AV (apical vesicles) region, which is located below PM1 and delimited towards the shank by a line connecting the distal endpoints of this PM domain, and 2) the UIM (unspecified internal membranes) region underneath PM3 with a depth of ten pixels corresponding to the average height of the AV region. Automatically extracted background-corrected intensity values (I) for all pixels in each ROIs were employed to calculate GP-values according to the following formula:

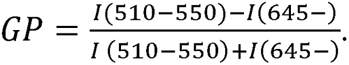

To determine the exact position and the width of the GP-value peak within the PM indicated by a dip in Di-4-ANEPPDHQ fluorescence emission in the upper (≥ 645 nm) channel, which was observed roughly between PM1 and PM2 (Fig. 3C), a search for a GP-value peak coinciding with this dip in fluorescence emission was performed within a PM region extending from 2.9 - 5.3 µm MD from the apex.

### Identification and sequence analysis of tobacco NtGEF12 isoforms

The *Nicotiana tabacum* genome databases TN90 and K326 (Sierro et al. 2014; https://solgenomics.net/) were BLAST searched using the full-length AtROPGEF12 amino acid sequence as query for proteins meeting the following criteria: at least 50 % amino acid identity shared with AtROPGEF12, annotated as AtROPGEF12-like protein, containing a full-length PRONE domain flanked by N- and C-terminal extensions, and presence of conserved target sites for RLK-mediated phosphorylation within the C-terminal extension. These searches resulted in the identification of three tobacco NtGEF12 isoforms, which are also represented in the Uniprot and NCBI databases: NtGEF12a (XP_016460052.1), NtGEF12b (XP_016479164.1) and NtGEF12c (XP_016503592.1). PRONE domains in these proteins were identified using the SMART tool (Letunic et al. 2021; Letunic and Bork 2018). Jalview 2.11.2.7 (Waterhouse et al. 2009) was employed to display and illustrate a Clustal Omega alignment (Madeira et al. 2022) of the full-length AtROPGEF12 and NtGEF12 amino acid sequences, and to generate a phylogenetic tree based on this alignment using the neighbor joining method and a BLOSUM62 substitution matrix.

### gDNA/RNA isolation, cDNA synthesis, RT-PCR and qRT-PCR

gDNA was isolated from mature tobacco leaves using the DNeasy^®^ plant Pro Kit (Qiagen). The TRIzol^TM^ reagent (Invitrogen) was employed to purify total RNA from mature flowers, stems and leaves of tobacco plants grown in soil, from roots of 7 d-old axenically cultured tobacco seedlings and from tobacco pollen tubes grown for 4 h in liquid PTNT medium. cDNA was synthesized using 1 µg of each total RNA preparation as template and the iScript^TM^ Reverse Transcription Supermix for qRT-PCR Kit (Bio-Rad). Primers listed in supplemental table S7 were employed for RT-PCR and qRT-PCR analysis of *NtGEF12* transcript levels in comparison to reference gene expression. To investigate gene expression patterns by RT-PCR, equal volumes of each cDNA sample and of the gDNA preparation were employed as PCR template. Fragments amplified using the FastGene Optima HotStart Ready Mix (Nippon Genetics) over 35 cycles in a C1000 Thermal Cycler (Bio-Rad) were subjected to agarose gel electrophoresis. The ubiquitously expressed tobacco *L25-RIBOSOMAL-PROTEIN* gene (Schmidt and Delaney 2010) was used as reference. qRT-PCR analysis of relative expression levels in pollen tubes was performed using pollen tube cDNA as template and the tobacco *E3-UBIQUITIN-LIGASE* gene (XM_016600104.1) as reference. The SsOAdvanced Universal SYBR Green Supermix (Bio-Rad) and a CFX96^TM^ Real-Time Cylcer (Bio-Rad) were employed for fragment amplification. Obtained fragments were verified by sequencing (Eurofins). Data were processed according to the 2^−ΔΔCt^ method (Livak and Schmittgen 2001).

### Mating-based split-ubiquitin assay

The haploid yeast strains THY.AP4 and THY.AP5 (Obrdiik et al. 2004) were transformed with pMetOYC-NtGEF12a or pNubG-NtRAC5, respectively, and plated on appropriate selective media (CSM +Ade, His, Trp, Ura or CSM +Ade, His, Leu; MP Biomedicals) according to Grefen et al. (2009). Instead of pNubG-NtRAC5, plasmids conferring NubG or NubWT expression were also introduced into THY.AP5 cells to generate control strains. 15 - 20 colonies transformed with each plasmid were used to inoculate selective liquid cultures, which were incubated overnight at 28°C under continuous shaking, before cells were pelleted and re-suspended in YPD medium (Roth). To allow mating, 5 µl aliquots of suspensions containing equal amounts of THY.AP4/pMetOYC-NtGEF12a cells and of either THY-AP5/pNubG-RAC5 cells, or of cells from control strains, were dropped onto YPD plates and incubated at 28°C for at least 6 h.

Colonies were then transferred to selective CSM +Ade, His (MP Biomedicals) plates, incubated at 28°C overnight and used to inoculate 5 ml liquid CSM +Ade, His. After overnight incubation at 28°C under continuous shaking, diploids obtained using different mating combinations were harvested from 100 µl cell suspension, re-suspended in sterile water and serially diluted to an OD_600_ of 1.0, 0.1 or 0.01. Finally, 7 µl of each diluted suspension were dropped onto CSM +Ade, His control plates and on selective CSM plates containing increasing amounts of methionine (Met; 0.5, 5, and 50 µM). Plates were incubated at 28°C and images were taken after 72 h.

Western blots were performed to confirm expression of OST-NtGEF12a-Cub-LexA-VP16 and NubG-2xHA-NtRAC5 fusion proteins in analyzed yeast cells. Total proteins were extracted using a post-alkaline procedure (Kushnirov 2000). OST-NtGEF12a-Cub-LexA-VP16 detection was performed using an anti-VP16 antibody (1:1,000; GeneTex, Cat. GTX30776) as primary antibody and anti-rabbit-POD (1:10,000; Merck-Millipore, Cat. AP307P) as secondary antibody. To detect NubG-2xHA-NtRAC5, a peroxidase-coupled anti-HA antibody (1:10,000; Roche, Cat. 12013819001) was employed.

### GenBank accession numbers

eYFP: AAO48591.1; NtGAP1: DQ813657; NtRISAP: AHX26274; NtINT4: XP_016480732; NtPI4P-5K4: XP_009802995.1; NtPLC3: EF043044; NtRAC5: AJ250174.1; NtGEF12a: XP_016460052.1; NtGEF12b: XP_016479164.1; NtGEF12c: XP_016503592.1; NtE3- UBIQUITIN-LIGASE: XM_016600104.1; NtL25-RIBOSOMAL-PROTEIN: L18908.

## Acknowledgments

The authors would like to thank Vanessa Schmidt and Martin Schuster for technical support, as well as Christina Müdsam for extracting and organizing data for GP-analysis. They are also grateful to Ingo Heilmann (cDNA encoding NtPI4P-5K4), Teun Munnik (cDNA encoding the PI4P marker eYFP-PH_FAPP1_), Martin Potocký (cDNA encoding the PA marker eYFP-Spo20p), Yvon Jaillais (cDNA encoding the PS marker eYFP-C2_Lact_), and Kai Konrad (transgenic tobacco line displaying expression of the Ca^2+^ marker 2xR-GECO1 in pollen tubes) for providing research materials.

## Author contributions

CF performed most experiments. JK (qRT-PCR) and CK (split-ubiquitin yeast two-hybrid assay) contributed specific data sets. SS generated and characterized transgenic tobacco lines. JA assisted in Di-4-ANEPPDHQ imaging, performed image deconvolution and developed software to analyze membrane order. CF, JK, CK, SM, IP and BK designed experiments. CF, TR, IP, SM and BK were responsible for data analysis and interpretation. CF, TR, IP and BK prepared a draft manuscript. CF and BK wrote the final version. The study was conceived by CF, IP and BK, and administered by BK. Funding was acquired by CF, IP, SM and BK.

## Supplemental data

The following materials are available in the online version of this article.

**Supplemental Figure S1:** Mean growth rates of all analyzed pollen tubes.

**Supplemental Figure S2:** Variable extension of the NtRAC5 domain at the PM of individual pollen tubes.

**Supplemental Figure S3:** Identification of tobacco homologs of AtROPGEF12.

**Supplemental Figure S4:** NtGEF12a interacts with NtRAC5 in split-ubiquitin yeast two-hybrid assays.

**Supplemental Figure S5:** Variable intracellular PS distribution in individual pollen tubes.

**Supplemental Figure S6:** Schematic display of the hardware employed to investigate membrane order based on widefield fluorescence microscopy.

**Supplemental Table S1:** Plant expression vectors used in this study.

**Supplemental Table S2:** Plasmids serving as templates to construct new expression cassettes.

**Supplemental Table S3:** Primers used to construct new expression cassettes or plasmids for split-ubiquitin assays.

**Supplemental Table S4:** Transgenic tobacco lines and imaging parameters used in this study.

**Supplemental Table S5:** MDs from the apex of maximal and half-maximal PM-associated relative fluorescence intensities (RFIs) displayed by all investigated PM domains or by free eYFP used as control.

**Supplemental Table S6:** Mean MDs from the apex of proximal and distal endpoints of all investigated PM domains.

**Supplemental Table S7:** Primers used for RT-PCR and qRT-PCR.

