## Supplemental Figures for "Plasma membrane and cytoplasmic compartmentalization: a dynamic structural framework required for pollen tube tip growth"

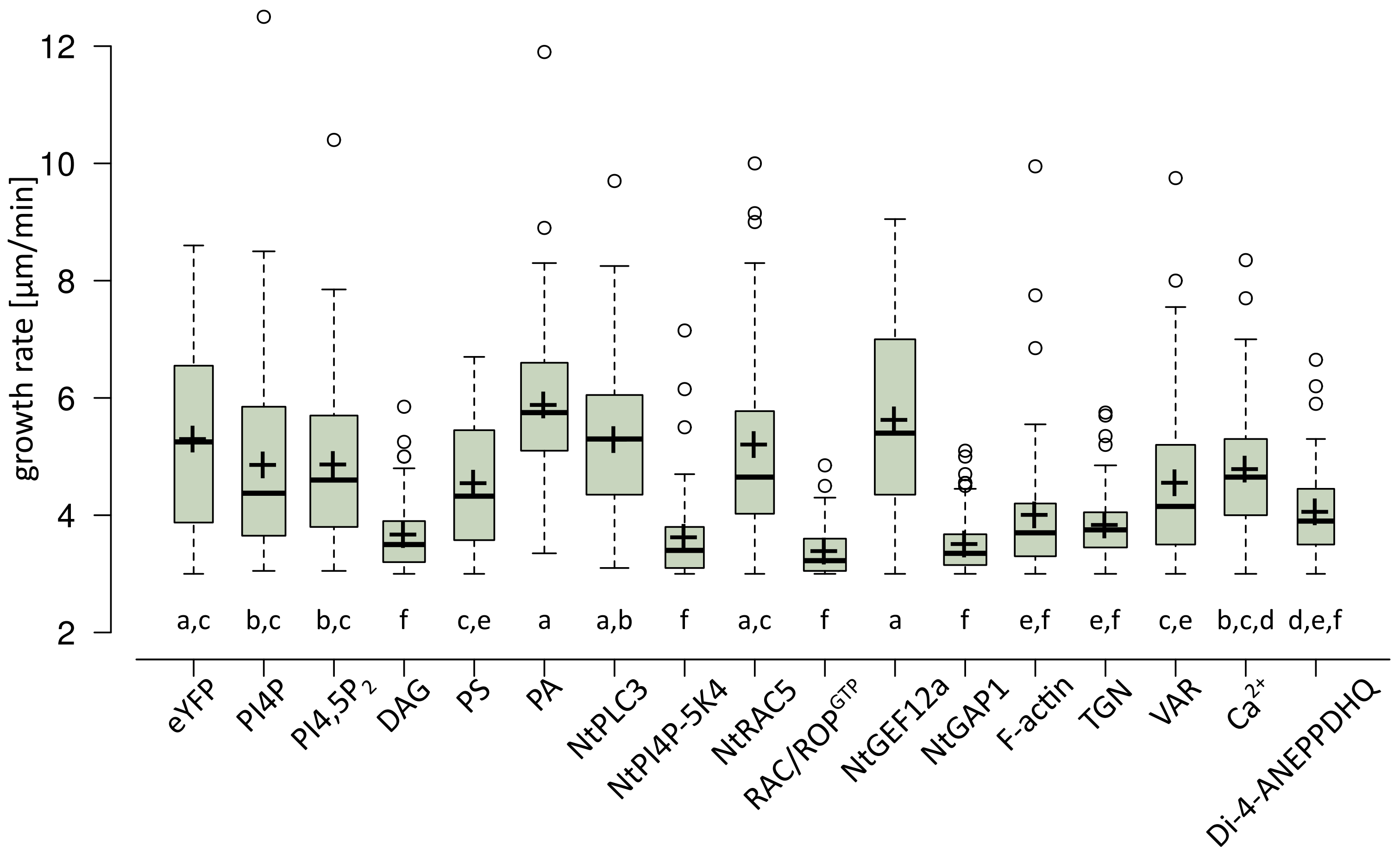


**Supplemental Figure S1: Mean growth rates of all analyzed pollen tubes.** Chart displaying the mean growth rates of all pollen tubes analyzed to generate the data shown in figures 2, 4, 6A and 7, which expressed a) free eYFP used as control, b) eYFP fusion proteins serving as markers for the indicated lipids, proteins or cytoplasmic structures and regions, or c) the $Ca^{2+}$ sensor 2xR-GECO1. Growth rates of individual pollen tubes were measured after recording medial confocal optical sections employed to characterize eYFP intracellular distribution (Fig. 2A) or to determine mean MDs from the apex of PM domain endpoints (Fig. 7A). A random selection of the same confocal optical sections was used to investigate relative fluorescence intensities (RFIs) associated with the PM (Figs. 2, 4, 6A and 7B). In addition, the chart displays the mean growth rate of all Di-4-ANEPPDHQ-labeled pollen tubes analyzed to determine membrane order (Fig. 3). Individual growth rates of these pollen tube were measured after recording widefield Z-stacks required for this purpose. Pollen tubes displaying growth rates of less than 3.0 µm/min were not included in any of the analyses outlined above. Datasets are displayed as boxplots indicating median (center line), mean (plus), upper and lower quartiles (box), minimum/maximum (whiskers) and outliers (dots). Box width is proportional to the square root of the number of analyzed pollen tubes: eYFP, n = 55 (see legend of figure 2A); PI4P, n = 70; $PI4,5P_2$, n = 83; DAG, n = 65; PS, n = 60; PA, n = 81; NtPLC3, n = 117; NtPI4P-5K4, n = 55; NtRAC5, n = 55; $RAC/ROP^{GTP}$, n = 66; NtGEF12a, n = 61; NtGAP1, n = 67; F-actin, n = 63; TGN, n = 69; VAR, n = 57; $Ca^{2+}$, n = 67 (see legend of figure 7A); Di-4-ANEPPDHQ, n = 50 (see legend of figure 3). Data were statistically analyzed using one-way ANOVA with Tukey Kramer (HSD) testing. Distinct letters indicate significant differences between data sets ($p \leq 0.05$).

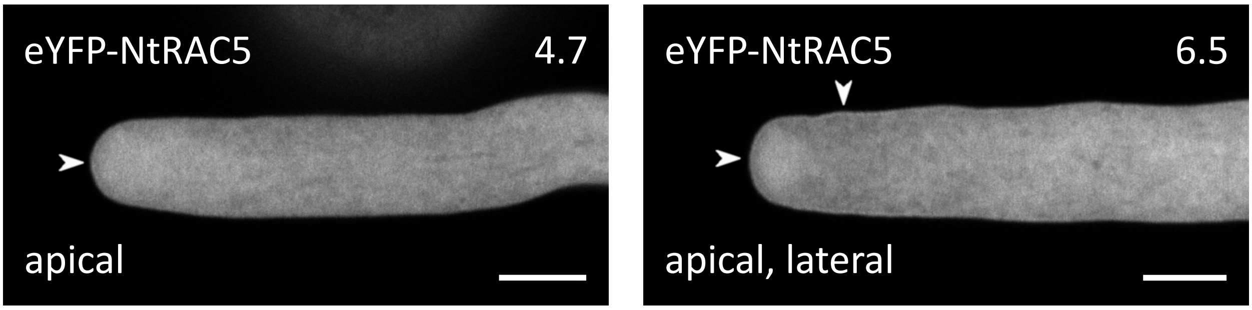


**Supplemental Figure S2: Variable extension of the NtRAC5 domain at the PM of individual pollen tubes.** Medial confocal optical sections through two different genetically identical pollen tubes stably expressing eYFP-NtRAC5 at minimal detectable levels, which display weak PM association of this fusion protein (arrowheads). In rare cases, eYFP-NtRAC5 PM association was detectable in such pollen tubes either exclusively at the extreme tip (left image) or within substantially larger apical and lateral regions (right image). Compared to all other PM domains analyzed in this study, the eYFP-NtRAC5 domain was exceptionally variable in extension. Numbers (top right) indicate the growth rates (µm/min) of the two pollen tubes shown, which were determined after image acquisition (average growth rate of all analyzed pollen tubes: Supplemental Fig. S1). Scale bars: 8 µm.

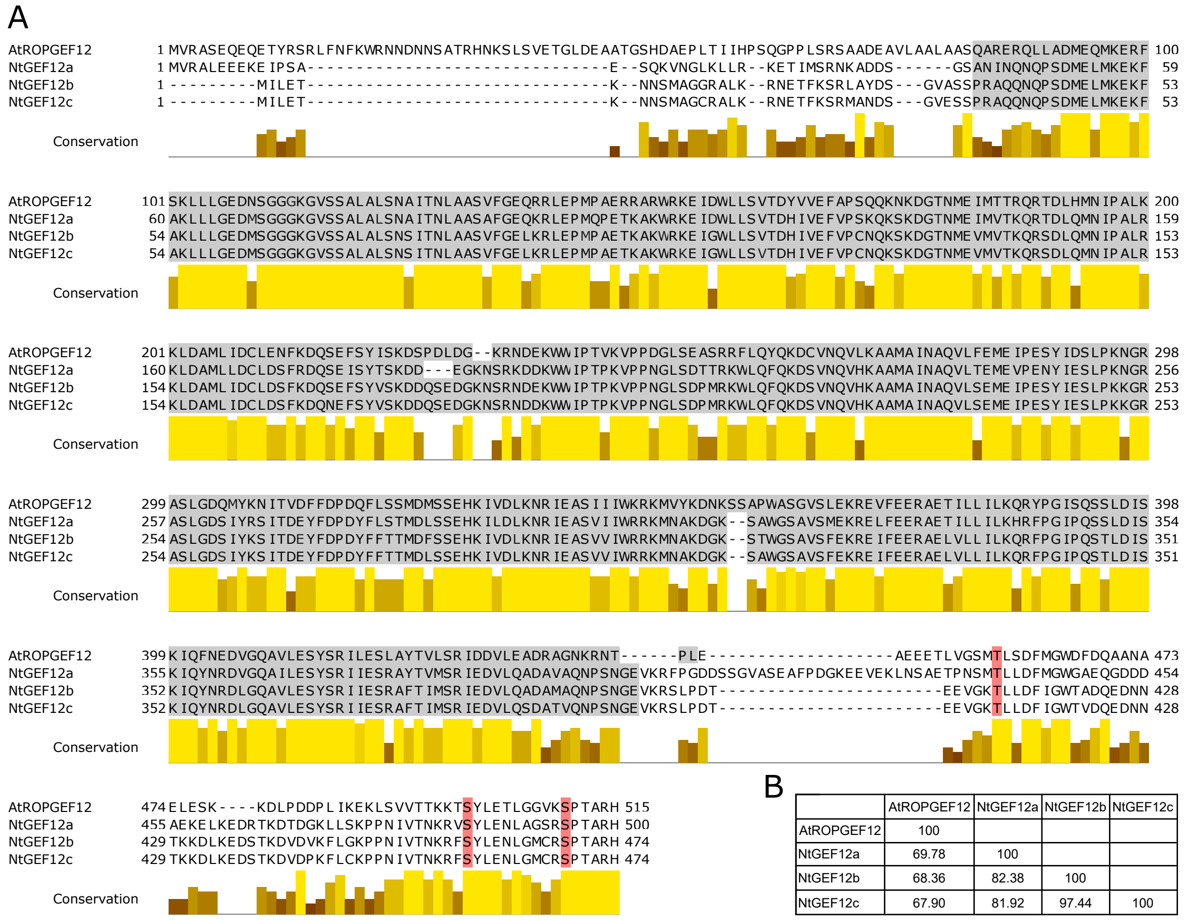


B

| | AtROPGEF12 | NtGEF12a | NtGEF12b | NtGEF12c |
|---|---|---|---|---|
| AtROPGEF12 | 100 | | | |
| NtGEF12a | 69.78 | 100 | | |
| NtGEF12b | 68.36 | 82.38 | 100 | |
| NtGEF12c | 67.90 | 81.92 | 97.44 | 100 |

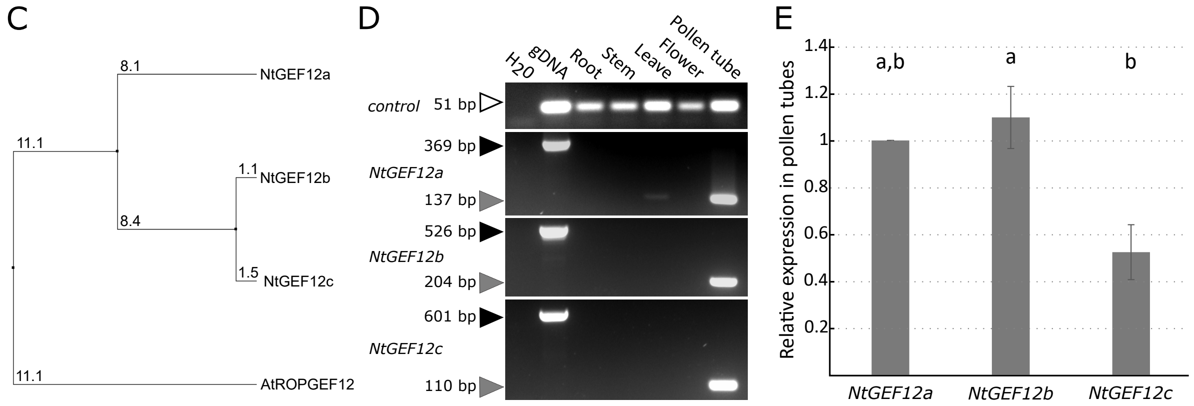


**Supplemental Figure S3: Identification of tobacco homologs of AtROPGEF12.** Full-length regions encoding three close AtROPGEF12 homologs were identified in available tobacco genome databases, which were designated NtGEF12a, NtGEF12b or NtGEF12c. **A)** Clustal Omega alignment of the amino acid sequences of AtROPGEF12 and of the identified tobacco homologs of this protein. Yellow bars: degree of conservation (taller bars and lighter color indicate higher conservation). Grey shading: highly conserved PRONE domain responsible for nucleotide exchange activity (annotation by SMART prediction). Red shading: Conserved S/T residues implicated in phosphoregulation of autoinhibited nucleotide exchange activity (Zhang and McCormick 2007).
**B)** Amino acid identities between the indicated proteins based on the alignment shown in A). **C)** Unrooted tree displaying phylogenetic relationships between the indicated proteins, which were inferred from the alignment of full-length amino acid sequences shown in A) (neighbor joining method). Numbers indicate percentage of non-identical amino acids. **D)** Semi-quantitative RT-PCR analysis of *NtGEF12a*, *b* and *c* transcript levels in different tobacco tissues. Intron spanning fragments were amplified to distinguish between cDNA (grey arrowheads) and genomic (gDNA, black arrowheads) templates. A fragment of a single exon of the gene encoding the ubiquitously expressed L25-RIBOSOMAL-PROTEIN was amplified as control (open arrowhead). **E)** qRT-PCR analysis of relative *NtGEF12a*, *b* and *c* transcript levels ($2^{-\Delta\Delta Ct}$ method; 6 biological replicates, each technically replicated trice), which confirms expression of all analyzed genes at similar levels in tobacco pollen tubes. The tobacco *E3-UBIQUITIN-LIGASE* reference gene and *NtGEF12a* transcript levels were used for data normalization. Data were statistically analyzed using one-way ANOVA with Tukey Kramer (HSD) testing. Distinct letters indicate significant differences between data sets ($p \leq 0.05$). Error bars: standard deviation.

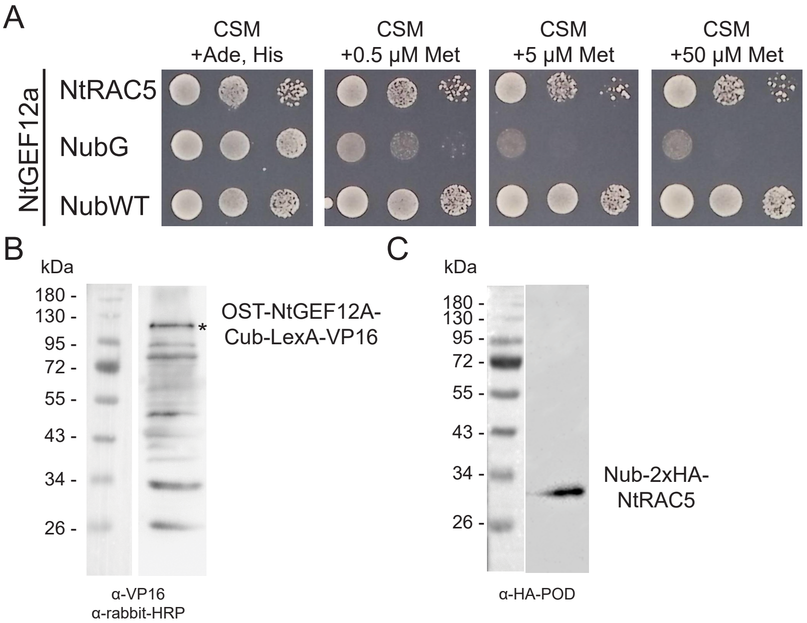


**Supplemental Figure S4: NtGEF12a interacts with NtRAC5 in split-ubiquitin yeast two-hybrid assays.**
**A)** Interaction between OST-NtGEF12a-Cub-LexA-VP16 (NtGEF12a) and NubG-2xHA-NtRAC5 (NtRAC5) at the yeast PM activated reporter gene expression restoring histidine and adenine biosynthesis. Free NubG and NubWT were employed as negative and positive controls, respectively. Serially diluted (left to right: 1.0, 0.1, and 0.01 $A_{600}$) yeast diploids were spotted on defined medium containing histidine and adenine (CSM +Ade, His) to verify mating, and on histidine/adenine-free medium containing methionine at the indicated concentrations (CSM +Met) to assay for interaction. **B)**, **C)** Verification of OST-NtGEF12a-Cub-LexA-VP16 (116.9 kDa) and NubG-2xHA-NtRAC5 (30 kDa) expression in analyzed yeast cultures based on immunoblotting. B) Anti-VP16 (primary) and anti-rabbit-HRP (secondary, conjugated to horseradish peroxidase) antibodies were employed to detect OST-NtGEF12a-Cub-LexA-VP16. C) NubG-2xHA-NtRAC5 was detected using an anti-HA-POD antibody (conjugated to peroxidase).

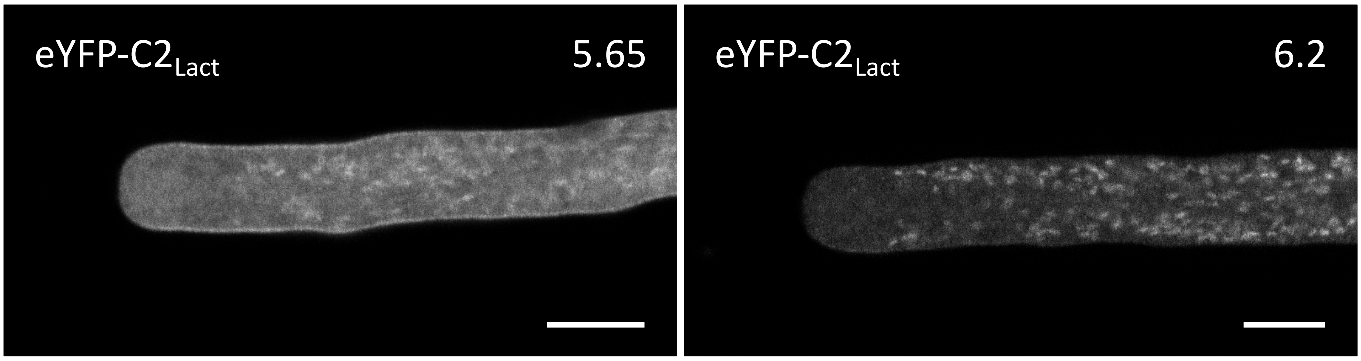


**Supplemental Figure S5: Variable intracellular PS distribution in individual pollen tubes.** Medial confocal optical sections through two different genetically identical pollen tubes, which stably expressed the PS marker eYFP-$C2_{Lact}$. Individual pollen tubes displayed variable relative levels of PS accumulation in the lateral PM, in apical vesicles and in other cytoplasmic endomembrane compartments (compare Fig. 6A and B [DMSO]). In rare cases, PS was barely detectable in the PM or in apical vesicles and strongly accumulated in unidentified cytoplasmic organelles (right image). Importantly, despite these variations in the observed PS distribution, this lipid was never detected in the PM at the apex of tobacco pollen tubes. Numbers (top right) indicate the growth rates (µm/min) of the two pollen tubes shown, which were determined after image acquisition (average growth rate of all analyzed pollen tubes: Supplemental Fig. S1). Scale bars: 8 µm.

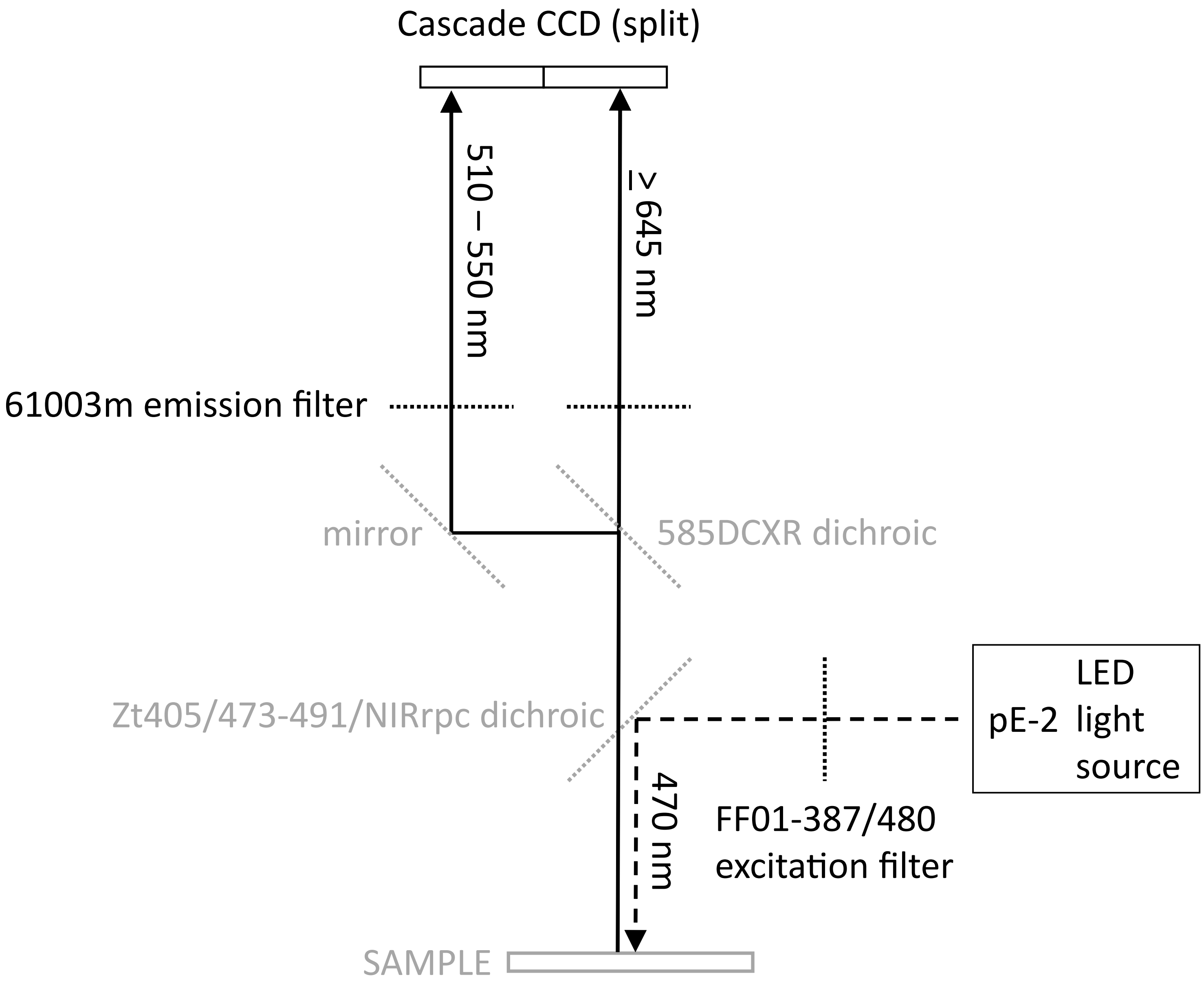


**Supplemental Figure S6: Schematic display of the hardware employed to investigate membrane order based on widefield fluorescence microscopy.** See material and methods for further information and a description of the imaging procedure.
