## Supplemental Tables for "Plasma membrane and cytoplasmic compartmentalization: a dynamic structural framework required for pollen tube tip growth"

**Supplemental Table S1: Plant expression vectors used in this study.** pHD342 was employed for transient pollen tube transformation. Expression cassettes from all other listed plasmids were subcloned into the binary vector pHD71 (Fritz and Kost 2020), which is derived from pPZP212 (Hajdukiewicz et al. 1994), for stable plant transformation. Templates, primers, target vectors and restriction enzymes used for PCR-based construction of new expression cassettes are indicated.

| Plasmid name | Expression cassette | DNA template[1] | Primer combination[2] | Target vector[3] | Restriction enzymes | Reference |
|---|---|---|---|---|---|---|
| pWEN240 | p*Lat52*:*eYFP*-*Nos*T | | | | | Klahre et al. 2006 |
| pFAU422 | p*Lat52*:*eYFP*-5xGA-*NtGEF12a*-*Nos*T | tobacco pollen tube cDNA | FAU-B122, FAU-B123 | pWEN240 | *Sal*I, *Xma*I | This study |
| pHD341 | p*Lat52*:*eYFP*-5xGA-*NtGAP1*-*Nos*T | | | | | Klahre and Kost 2006 |
| pHD342 | p*Lat52*:$CRIB_{NtGAP1}$-5xGA-*eYFP*-*Nos*T | | | | | Klahre and Kost 2006 |
| pHD23 | p*Lat52*:*eYFP*-5xGA-*NtRAC5*-*Nos*T | | | | | Sun et al. 2015 |
| pFAU581 | p*Lat52*:*NtPI4P-5K4*-5xGA-*eYFP*-*Nos*T | pSLUobt13 | FAU-B910, FAU-B911 | pHD32 | *Ngo*MIV, *Apa*I | This study |
| pHD133 | p*Lat52*:*NtPLC3*-*eYFP*-*Nos*T | | | | | Helling et al. 2006 |
| pFAU699 | p*Lat52*:*eYFP*-5xGA-$PH_{FAPP1}$-*Nos*T[4] | pFAUobt100 | FAU-C459, FAU-C460 | pWEN240 | *Ngo*MIV, *Apa*I | This study |
| pFAU780 | p*Lat52*:*eYFP*-5xGA-$C2_{Lact}$-*Nos*T[5] | pFAUobt104 | FAU-C624, FAU-C625 | pWEN240 | *Ngo*MIV, *Sac*II | This study |
| pWEN106 | p*Lat52*:*eYFP*-$PH_{PLC\delta 1}$-*Nos*T[6] | | | | | Helling et al. 2006 |
| pFAU131 | p*Lat52*:*eYFP*-5xGA-*Spo20p*-*Nos*T[7] | | | | | Potocký et al. 2014 |
| pHD274 | p*Lat52*:$C1_{PKC\gamma}$-*eYFP*-*Nos*T[8] | | | | | Helling et al. 2006 |
| pFAU351 | p*Lat52*:*NtINT4*-*eYFP*-*Nos*T | | | | | Grebnev et al. 2020 |
| pHD86 | p*Lat52*:*eYFP*-5xGA-*NtRISAP*-*Nos*T | | | | | Stephan et al. 2014 |
| pSLU29 | p*Lat52*:*LIFEACT*-5xGA-*eYFP*-*Nos*T | | | | | Montes-Rodriguez and Kost 2017 |

[1]see supplemental table S2 for further information; [2]see supplemental table S3 for primer sequences; [3]Klahre and Kost 2006; Grebnev et al. 2020; [4]contains the PLECKSTRIN HOMOLOGY (PH) domain of human PHOSPHATIDYL-INOSITOL-4-PHOSPHATE ADAPTOR PROTEIN-1 (FAPP1); [5]contains the PS-binding C2 domain of bovine LACTADHERIN (Lact); [6]contains the PH domain of human PHOSPHOLIPASE Cδ1 (PLCδ1); [7]contains the PA-binding domain of the yeast SNARE protein (Spo20p); [8]contains the C-terminal end of the first cysteine-rich domain (C1) of rat PROTEIN KINASE Cγ (PKCγ).

**Supplemental Table S2: Plasmids serving as templates to construct new expression cassettes (see Supplemental Table S1).**

| Plasmid name | Original name/description | Reference |
|---|---|---|
| pSLUobt13[1] | p*LAT52:464-YFP* (containing *NtPI4P-5K4* cDNA) | XP_009802995.1 |
| pFAUobt100 | p*Nigel07-PH*$_{FAPP1}$ containing p*UBQ10*:*eYFP-PH*$_{FAPP1}$ cassette | Thole et al. 2008 |
| pFAUobt104 | *UBQ10*prom:*mCITRINE*noSTOP-*LactC2*/pH7m34GW | Platre et al. 2018 |

[1]obtained from Heilmann Lab, Martin-Luther-University of Halle-Wittenberg. The *NtPI4P-5K4* cDNA was amplified from a tobacco pollen tube cDNA library (Klahre et al. 2006) using degenerate primers as described by Stenzel et al. (2012).

**Supplemental Table S3: Primers used to construct expression cassettes or plasmids for split-ubiquitin assays.** Protection bases: bold. Restriction enzyme recognition sites: underlined. Stop codons: italics.

| Primer name | 5'-3'-nucleotide sequence | Description |
|---|---|---|
| FAU-B122 | **TAT**GTCGACATGGTTCGAGCACTAGAAGAA | Protection bases, *Sal*I site<br>fw; Amplification of *NtGEF12a* |
| FAU-B123 | **TAT**CCCGGGGTGGCGTGCTGTTGGAC | Protection bases, *Xma*I site<br>rev; Amplification of *NtGEF12a* |
| FAU-B910 | GCCGGCATGGAAAAGGCTTGGGAGG | *Ngo*MIV site<br>fw; Amplification of *NtPI4P-5K4* |
| FAU-B911 | CCCGGGAGTGTCCTCTGTAAAAACTTTGAA | *Apa*I site<br>rev; Amplification of *NtPI4P-5K4* |
| FAU-C459 | GCCGGCATGGAGGGGGTGTTGTACAAG | *Ngo*MIV site<br>fw; Amplification of $PH_{FAPP1}$ |
| FAU-C460 | GGGCCC*TTA*TGTATCAGTCAAACATGCTTTGG | *Apa*I site; Stop codon;<br>rev; Amplification of $PH_{FAPP1}$ |
| FAU-C624 | GCCGGCTGCACTGAACCCCTAGGCC | *Ngo*MIV site<br>fw; Amplification of $C2_{Lact}$ |
| FAU-C625 | CCGCGGC*TAA*CAGCCCAGCAGCTCC | *Sac*II site; Stop codon;<br>rev; Amplification of $C2_{Lact}$ |
| SU-1 | **TAT**GGATCCTATGGTTCGAGCACTAGAAGAAGA | Protection bases, *Bam*HI site<br>fw; Amplification of *NtGEF12a*<br>> pMetOYC-NtGEF12a |
| SU-2 | **ATT**CTCGAGTTGTGGCGTGCTGTTGGACTTCT | Protection bases, *Xho*I site<br>rev; Amplification of *NtGEF12a*<br>> pMetOYC-NtGEF12a |
| SU-3 | **TTA**GGATCCATGAGTGCTTCAAGGTTTATC | Protection bases, *Bam*HI site<br>fw; Amplification of *NtRAC5*<br>> pNubG-NtRAC5 |
| SU-4 | **AAT**CTCGAG*TCA*CAATATCGAGCAGGATT | Protection bases, *Xho*I site; Stop codon<br>rev; Amplification of *NtRAC5*<br>> pNubG-NtRAC5 |

**Supplemental Table S4: Transgenic tobacco lines and imaging parameters used in this study.**

| Fusion protein expressed | Line identifier | Lightning mode | Scan speed | Line averaging | Dynamic range | Resolution | Reference |
|---|---|---|---|---|---|---|---|
| eYFP-$PH_{FAPP1}$ (PI4P) | FAU777 11 | no | 400 Hz | 3 | 8 bit | 1024x1024 | This study |
| eYFP-$PH_{PLC\delta 1}$ ($PI4,5P_2$) | FAU694 9_6; 9_12; 9_14 | no | 400 Hz | 3 | 8 bit | 1024x1024 | This study |
| $C1_{PKC\gamma}$-eYFP (DAG) | 4_13 | yes | 598 Hz | 3 | 16 bit | 1424x1424 | This study |
| eYFP-$C2_{Lact}$ (PS) | FAU790 7; 7_2; 7_4 | no | 400 Hz | 3 | 8 bit | 1024x1024 | This study |
| eYFP-Spo20p (PA) | 12_12 | no | 400 Hz | 3 | 8 bit | 1024x1024 | This study |
| NtPLC3-eYFP | 11_11 | no | 600 Hz | 3 | 8 bit | 1024x1024 | This study |
| NtPI4P5-K4-eYFP | FAU682 6_1 | yes | 700 Hz | 2 | 16 bit | 1280x1280 | This study |
| eYFP-NtRAC5 | 5_15 | no | 400-600 Hz | 5 | 8 or 16 bit | 1024x1024 | This study |
| $CRIB_{NtGAP1}$-eYFP ($RAC/ROP^{GTP}$) | - | no | 600 Hz | 3 | 8 bit | 1024x1024 | transgenic lines could not be established |
| eYFP-NtGEF12a | 1_15; 1_2 | no | 400 Hz | 3 | 8 bit | 1024x1024 | This study |
| eYFP-NtGAP1 | 15_12 | no | 600 Hz | 5 | 8 bit | 1024x1024 | This study |
| LIFEACT-eYFP (F-actin) | FAU162 1, 1_1; 4 | no | 400 Hz | 2 | 8 bit | 1024x1024 | Stephan et al. 2014 |
| eYFP-NtRISAP (TGN) | 71_86 | no | 400 Hz | 3 | 8 bit | 1024x1024 | Stephan et al. 2014 |
| NtINT4-eYFP (VAR) | FAU656 3_5 | no | 600 Hz | 3 | 8 bit | 1024x1024 | Grebnev et al. 2020 |
| 2xR-GECO1[1] ($Ca^{2+}$) | 8Nr8_3 | no | 400 Hz | 1 | 8 bit | 1024x1024 | Li et al. 2021 |
| eYFP | FAU778 13 | no | 600 Hz | 3 | 8 bit | 1024x1024 | This study |

[1]R-GECO1 comprises circularly permuted mApple fused a) at the N-terminus to the calmodulin (CaM)-binding region of chicken myosin light chain kinase (M13) and b) at the C-terminus to a vertebrate CaM (Zhao et al. 2011).

**Supplemental Table S5: MDs from the apex of maximal and half-maximal PM-associated relative fluorescence intensities (RFIs) displayed by all investigated PM domains or by free eYFP used as control.** The table lists mean MDs from the apex of maximal RFIs, as well as of proximal and distal half-maximal RFIs (if available), along with the distance between the proximal and distal half-maximal RFIs, if both these values are available. In case one of the half-maximal RFIs is not available, the distance between the maximal and the other half-maximal RFI is indicated.

| PM domain or free eYFP | MD from the apex [µm] | | | Distance between distal & proximal half-maximal RFI [µm] |
|---|---|---|---|---|
| | Proximal half-maximal RFI | Maximal RFI | Distal half-maximal RFI | |
| PI4P (eYFP-PH$_{FAPP1}$) | 2.381 | 4.04 | 7.287 | 4.906 |
| PI4,5P$_2$ (eYFP-PH$_{PLC\delta1}$) | 2.309 | 4.401 | 7.648 | 5.339 |
| DAG (C1$_{PKC\gamma}$-eYFP) | 2.02 | 4.329 | 11.616 | 9.596 |
| PS (eYFP-C2$_{Lact}$) | 5.123 | 9.884 | 22.871 | 17.748 |
| PA (eYFP-Spo20p) | 5.628 | 10.39 | 19.625 | 13.997 |
| NtPLC3 (NtPLC3-eYFP) | 5.916 | 13.781 | 25.036 | 19.12 |
| NtPI4P-5K4 (NtPI4P5K4-eYFP) | 4.978 | 8.441 | 12.554 | 7.576 |
| NtRAC5 (eYFP-NtRAC5) | 0 | 0.072 | 6.782 | 6.782 |
| RAC/ROP$^{GTP}$ (CRIB$_{NtGAP1}$-eYFP) | | 0 | 2.742 | 2.742 |
| NtGEF12a (eYFP-NtGEF12a) | 0 | 0.072 | 1.804 | 1.804 |
| NtGAP1 (eYFP-NtGAP1) | 3.752 | 5.988 | 11.039 | 7.287 |
| F-actin fringe (LIFEACT-eYFP) | 3.824 | 4.906 | 7.792 | 3.968 |
| TGN (eYFP-NtRISAP) | 2.597 | 4.257 | 6.061 | 3.464 |
| VAR (NtINT4-eYFP) | | 0 | 3.824 | 3.824 |
| $Ca^{2+}$ (2xR-GECO1) | 0.072 | 0.505 | 3.896 | 3.824 |
| eYFP | 22.944 | 53.319 | | 30.735 |

**Supplemental Table S6: Mean MDs from the apex of proximal and distal endpoints of all investigated PM domains.** Mean values and standard deviations are listed. Rounded values without standard deviations are indicated in figures 1 and 7A.

| PM domain | MD from the apex [μm] | | Domain length [μm] |
|---|---|---|---|
| | Proximal end | Distal end | |
| PI4P (eYFP-$PH_{FAPP1}$) | 0 | 9.552 ± 1.731 | 9.552 ± 1.731 |
| $PI4,5P_2$ (eYFP-$PH_{PLC\delta 1}$) | 0 | 10.129 ± 1.757 | 10.129 ± 1.757 |
| DAG ($C1_{PKC\gamma}$-eYFP) | 0 | 12.084 ± 2.392 | 12.084 ± 2.392 |
| PS (eYFP-$C2_{Lact}$) | 6.011 ± 1.080 | 28.669 ± 6.127 | 22.658 ± 6.088 |
| PA (eYFP-Spo20p) | 6.085 ± 0.831 | 21.342 ± 2.440 | 15.258 ± 2.399 |
| NtPLC3 (NtPLC3-eYFP) | 6.286 ± 0.942 | 27.229 ± 3.560 | 20.943 ± 3.477 |
| NtPI4P-5K4 (NtPI4P-5K4-eYFP) | 5.697 ± 1.374 | 13.496 ± 1.863 | 7.800 ± 1.704 |
| NtRAC5 (eYFP-NtRAC5) | 0 | 18.804 ± 7.123 | 18.804 ± 7.123 |
| $RAC/ROP^{GTP}$ ($CRIB_{NtGAP1}$-eYFP) | 0 | 3.271 ± 0.738 | 3.271 ± 0.738 |
| NtGEF12a (eYFP-NtGEF12a) | 0 | 2.998 ± 0.668 | 2.998 ± 0.668 |
| NtGAP1 (eYFP-NtGAP1) | 4.175 ± 0.751 | 11.178 ± 1.719 | 7.002 ± 1.812 |
| F-actin fringe (LIFEACT-eYFP) | 5.037 ± 0.659 | 7.070 ± 0.814 | 2.027 ± 0.529 |
| TGN (eYFP-NtRISAP) | 3.310 ± 0.834 | 6.647 ± 0.680 | 3.337 ± 0.653 |
| VAR (NtINT4-eYFP) | 0 | 4.455 ± 1.244 | 4.455 ± 1.244 |
| $Ca^{2+}$ (2xR-GECO1) | 0 | 5.845 ± 0.980 | 5.845 ± 0.980 |

**Supplemental Table S7: Primers used for RT-PCR and qRT-PCR.**

| Primer name | 5’-3’-nucleotide sequence | Description |
|---|---|---|
| FAU995 | GGACTTGGATCACCTGCT | *NtE3-UBIQUITIN-LIGASE*, fw reference gene, XM_016600104.1 |
| FAU996 | TGGACATACAACCTCATAACTCTG | *NtE3-UBIQUITIN-LIGASE*, rev reference gene, XM_016600104.1 |
| FAU-E287 | CCCCTCACCACAGAGTCTGC | *NtL25-RIBOSOMAL-PROTEIN*, fw reference gene, L18908 (Schmidt and Delaney 2010) |
| FAU-E288 | AAGGGTGTTGTTGTCCTCAATCTT | *NtL25-RIBOSOMAL-PROTEIN*, rev reference gene, L18908 (Schmidt and Delaney 2010) |
| FAU-E307 | GACGATTCAGGCTCCGCTAAC | *NtGEF12a*, Exon 1, fw XP_016460052.1, |
| FAU-E308 | GACAATGCCAATGCTGATGAAACA | *NtGEF12*a, Exon 2, rev XP_016460052.1 |
| FAU-E309 | GTCTCGTATGGCTAATGATTCAGGT | *NtGEF12c*, Exon 1, fw XP_016503592.1 |
| FAU-E310 | CGCCAAGGAGCAACTTTGC | *NtGEF12c*, Exon 2, rev XP_016503592.1 |
| FAU-E311 | CAATTCCATGGCTGGGGGC | *NtGEF12b*, Exon 1, fw XP_016479164.1 |
| FAU-E312 | AAAGCCAGCGCAGAAGAGAC | *NtGEF12b*, Exon 2, rev XP_016479164.1 |
